# ncBAF, a chromatin remodeler, enhances PXR-mediated transcriptional activation in the human and mouse liver

**DOI:** 10.1101/2023.02.03.527063

**Authors:** Kiamu Kurosawa, Masataka Nakano, Itsuki Yokoseki, Mai Nagaoka, Seiya Takemoto, Yoshiyuki Sakai, Kaoru Kobayashi, Yasuhiro Kazuki, Tatsuki Fukami, Miki Nakajima

## Abstract

Pregnane X receptor (PXR) is one of the key regulators of drug metabolism, gluconeogenesis, and lipid synthesis in the human liver. Activation of PXR by drugs such as rifampicin, simvastatin, and efavirenz causes adverse reactions such as drug□drug interaction, hyperglycemia, and dyslipidemia. The inhibition of PXR activation has merit in preventing such adverse events. Here, we demonstrated that bromodomain containing protein 9 (BRD9), a component of non-canonical brahma-related gene 1-associated factor (ncBAF), one of the chromatin remodelers, interacts with PXR. Rifampicin-mediated induction of CYP3A4 expression was attenuated by iBRD9, an inhibitor of BRD9, in human primary hepatocytes and CYP3A/PXR-humanized mice, indicating that BRD9 enhances the transcriptional activation of PXR *in vitro* and *in vivo*. Chromatin immunoprecipitation assay reveled that iBRD9 treatment resulted in attenuation of the rifampicin-mediated binding of PXR to the *CYP3A4* promoter region, suggesting that ncBAF functions to facilitate the binding of PXR to its response elements. Efavirenz-induced hepatic lipid accumulation was attenuated by iBRD9 in C57BL/6J mice, suggesting that the inhibition of BRD9 would be useful to reduce the risk of efavirenz-induced hepatic steatosis. Collectively, we found that inhibitors of BRD9, a component of ncBAF that plays a role in assisting transactivation by PXR, would be useful to reduce the risk of PXR-mediated adverse reactions.

## Introduction

Pregnane X receptor (PXR) is a nuclear receptor that plays important roles in the regulation of xenobiotic metabolism in the human liver and intestine. It is activated by xenobiotics such as rifampicin and ritonavir (1–2) and induces several drug-metabolizing enzymes such as cytochrome P450 (CYP) 3A4, CYP2C9, CYP2B6, and UDP-glucuronosyltransferase (UGT) 1A1 (3–4), potentially causing adverse drug□drug interactions. For example, the concomitant use of St. John’s wort containing hyperforin, a ligand of PXR, results in insufficient immunosuppressive effects of tacrolimus by induction of CYP3A4 catalyzing the metabolism of tacrolimus (5).

In addition to drug-metabolizing enzymes, PXR regulates the expression of gluconeogenic enzymes and lipid synthases (6). Previous clinical studies have reported that treatment with typical activators of PXR, such as simvastatin, increases blood glucose levels in patients (7–8). As a potential mechanism of drug-induced hyperglycemia, the induction of gluconeogenic enzymes such as glucose-6-phosphatase (G6Pase) and phosphoenolpyruvate carboxykinase 1 (PEPCK1) by activated PXR in the liver has been suggested (9). Efavirenz, a non-nucleoside reverse transcriptase inhibitor, is known to cause dyslipidemia (10). A recent study revealed that efavirenz-induced accumulation of triglyceride and cholesterol in the liver is due to the induction of lipogenic genes, including CD36 and squalene epoxidase (SQLE), by activated PXR (11). Preventing PXR activation would yield the benefit of avoiding adverse drug events, but no clinically practical inhibitors for PXR activation have been developed. Expanding our understanding of the regulatory mechanisms of PXR activation may provide a clue to avoid such adverse events.

Over the past two decades, the following molecular mechanisms for PXR-mediated transcriptional activation have been disclosed. In the transcriptionally inactive state, PXR forms a complex with heat shock protein 90 and constitutive androstane receptor retention protein in the cytoplasm (12). Upon activation by a ligand, PXR translocates into the nucleus and forms a heterodimer with retinoid X receptor α (RXRα). Then, the PXR-RXRα dimer binds to the response elements in the 5’-flanking regions of the target genes to activate their transcription (13–15). In addition, several PXR coactivators, such as steroid co-activator 1 and proliferator-activated receptor-γ coactivator 1α, bind to PXR-RXRα to promote PXR-mediated gene transcription (16–17).

In the present study, to uncover novel transregulators of PXR, we performed co-immunoprecipitation using anti-PXR antibody against human liver-derived cell lysate followed by proteomic mass spectrometric analysis. We noticed that PXR interacts with components of switch/sucrose non-fermentable (SWI/SNF) complexes in a ligand-dependent manner. SWI/SNF complexes are ATP-dependent chromatin remodelers (18). This chromatin remodeling complex comprises of more than 10 units including ATPase subunits, core subunits, and accessory subunits. SWI/SNF complexes are classified into three major members, canonical brahma-related gene 1-associated factor (cBAF), polybromo-associated BAF (PBAF), and non-canonical BAF (ncBAF) according to the included unique units (19). The main ATPase subunits such as brahma-related gene (BRG) 1 and brahma (BRM), and the core subunits such as BAF47, BAF155, and BAF170 are common to the three members (Fig. 1C). These units rearrange the chromatin structure between the condensed state and transcriptionally active state, thereby affecting gene expression (20). Specific accessory units such as AT-rich interactive domain-containing protein (ARID) 1A and double PHD fingers (DPF) 2 for cBAF; bromodomain containing protein (BRD) 7, ARID2 and polybromo (PBRM) 1 for PBAF; and BRD9 and glioma tumor suppressor candidate region gene 1 (GLTSCR1) /GLTSCR1-like (GLTSCR1L) for ncBAF are believed to contribute to specific targeting of SWI/SNF complexes to chromatin through interactions with transcription factors or histones (21). In particular, BRD7 and BRD9 are important units recognizing the acetylated lysine residues of transcription factors or histones via their bromodomain (22). Knowledge about the role of each SWI/SNF complex in cancer development has gradually accumulated (23), but its significance in hepatic function including drug metabolism, lipid metabolism, and glucose metabolism remains to be clarified. In this study, we aimed to clarify the role of the SWI/SNF complex in PXR-mediated transcriptional activation and to investigate whether the inhibition of SWI/SNF complex is useful for attenuating PXR-mediated drug□drug interaction and adverse reactions.

**Figure 1.**
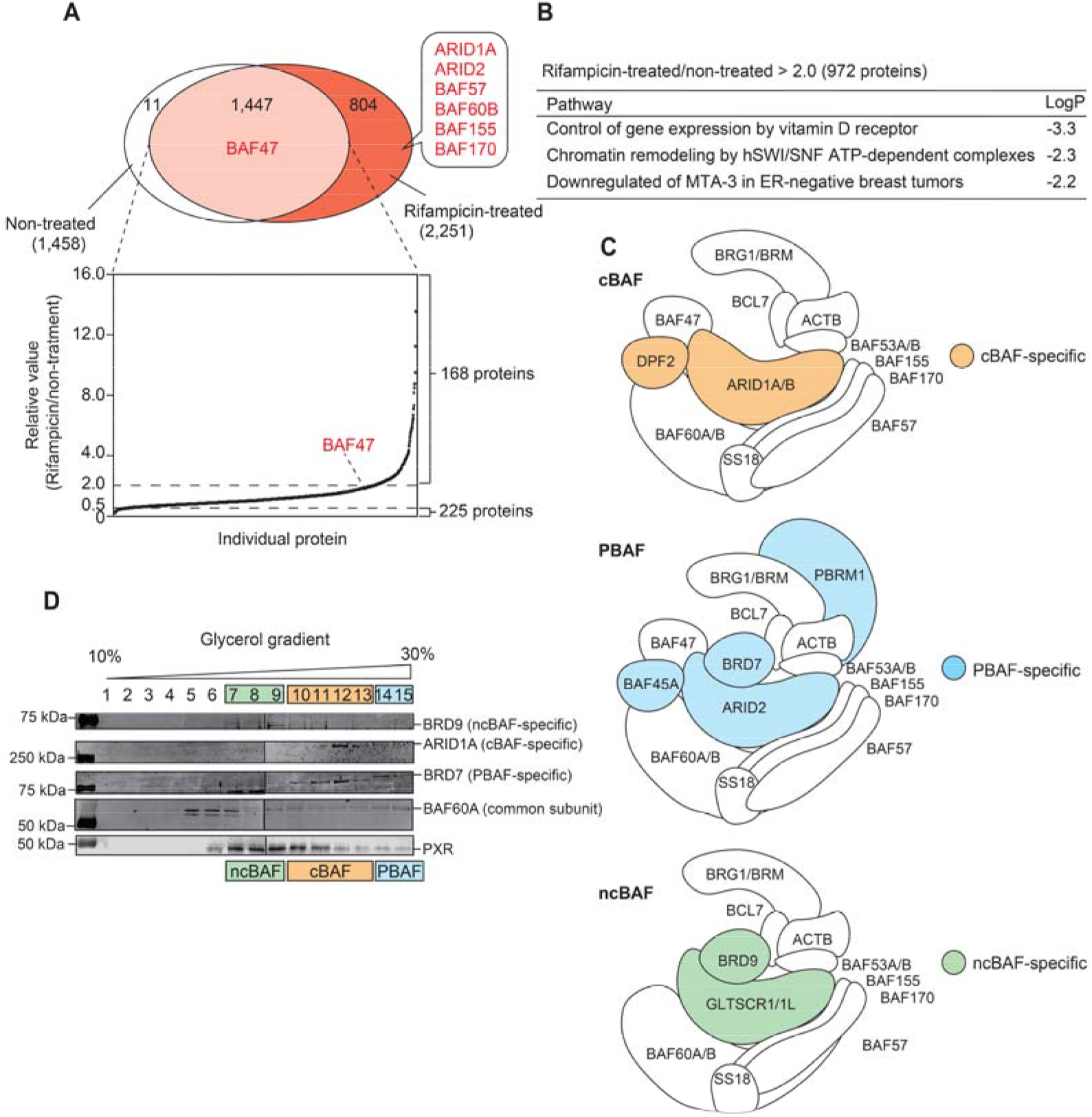
Identification of proteins interacting with PXR. ShP51 cells were treated with rifampicin for 24 hr. Co-immunoprecipitation with anti-PXR antibody was performed using whole cell lysates followed by mass spectrometric analysis. (A) The numbers of proteins detected in the immunoprecipitant of rifampicin-and/or non-treated cells are shown in a Venn diagram. The ratios of the protein amounts of the rifampicin-treated sample to those of non-treated sample are shown in the bottom graph. (B) The proteins whose abundance ratio rifampicin-treated/non-treated was > 2.0 (168 proteins) as well as proteins detected only in rifampicin-treated sample (804 proteins) were subjected to pathway analysis using DAVID Bioinformatics Resources (https://david.ncifcrf.gov/home.jsp). (C) Schematic representation of BAF, PBAF, and ncBAF composition reported by ref. 21 and ref. 58. (D) Glycerol gradient sedimentation was performed using ShP51 cell nuclear extracts, and ARID1A, BRD7, BRD9, and PXR proteins were detected by Western blotting. Source data are available online for this figure.

## Results

### Identification of proteins that interact with PXR in ShP51 cells

To identify proteins interacting with PXR in the process of PXR activation, we performed co-immunoprecipitation using lysates from rifampicin-treated and non-treated ShP51 cells followed by mass spectrometric analysis. As a result, 2,251 and 1,458 proteins were detected in the rifampicin-treated and non-treated samples, respectively. Among them, 1,447 proteins were common to both samples. Among the commonly detected proteins, 168 and 225 proteins showed more than 2.0-fold higher and less than 0.5-fold lower ratios of abundance in rifampicin-treated samples than in non-treated samples, respectively (Fig. 1A). Considering the possibility that the immunoprecipitated proteins may affect PXR-mediated transcriptional activation, we performed a pathway analysis targeting proteins whose abundance ratio rifampicin-treated/non-treated was > 2.0 (168 proteins) as well as proteins detected only in rifampicin-treated sample (804 proteins) using DAVID Bioinformatics Resources (https://david.ncifcrf.gov/home.jsp). As a result, three pathways, 1) control of gene expression by vitamin D receptor (VDR) pathway, 2) chromatin remodeling by hSWI/SNF ATP-dependent complexes pathway, and 3) downregulated of MTA-3 in ER-negative breast tumors pathway (Fig. 1B) were identified with statistical significance. SWI/SNF complexes are required for transcriptional activation by nuclear receptors such as estrogen receptor, progesterone receptor, androgen receptor (AR), glucocorticoid receptor (GR), peroxisome proliferator-activated receptor γ, and VDR (24), but significant roles of chromatin remodelers in PXR-mediated transcriptional regulation remain to be determined. The proteomic analysis showed that the interactions of PXR with BAF57, BAF47, BAF170, BAF155, and BAF60B, common subunits of SWI/SNF complexes, ARID1A, a cBAF-specific subunit, and ARID2, a PBAF-specific subunit, were increased by rifampicin treatment (Fig. 1A and 1C). The molecular weights of cBAF, PBAF, and ncBAF were calculated as 1.1, 1.4, and 0.9 MDa based on their constituents, respectively, and each complex can be separated by glycerol gradient sedimentation (21). To determine the SWI/SNF sub-complexes which interact with PXR, glycerol gradient sedimentation for nuclear extracts prepared from ShP51 cells was conducted, followed by Western blotting. As the results, BRD9, ARID1A, BRD7, and BAF60A were detected in fractions 7-9, 10-13, 14-15, and 7-15, respectively, and the order was consistent with the study reported by ref. 21. PXR was mainly detected in fractions 7-9, indicating that PXR interacts with ncBAF (Fig. 1D). In a subsequent study, we sought to clarify the functional significance of chromatin remodeling by ncBAF in PXR-mediated transcriptional regulation.

### PXR interacts with BRD9

BRD7 and BRD9 are known to recognize acetylated lysine residues of transcription factors via their bromodomain (22). It has been reported that the lysine 109 (K109) in the T-box domain of PXR is acetylated (25). Considering the possibility that the interaction of PXR with ncBAF complex but not PBAF (shown in Fig. 1D) might be due to the difference in the recognition of BRD9 but not BRD7 against PXR, we performed co-immunoprecipitation followed by Western blotting. As shown in Fig. 2A, the band intensity of BRD9-His was higher than that of BRD7-His in the immunoprecipitant using anti-FLAG antibody, indicating that BRD9 potently interacts with PXR. To investigate whether PXR directly interacts with BRD9, pull-down assay was performed. Since the same results as Fig. 2A was observed (Fig. S1A), it was suggested that PXR directly interacts with BRD9.

**Figure 2.**
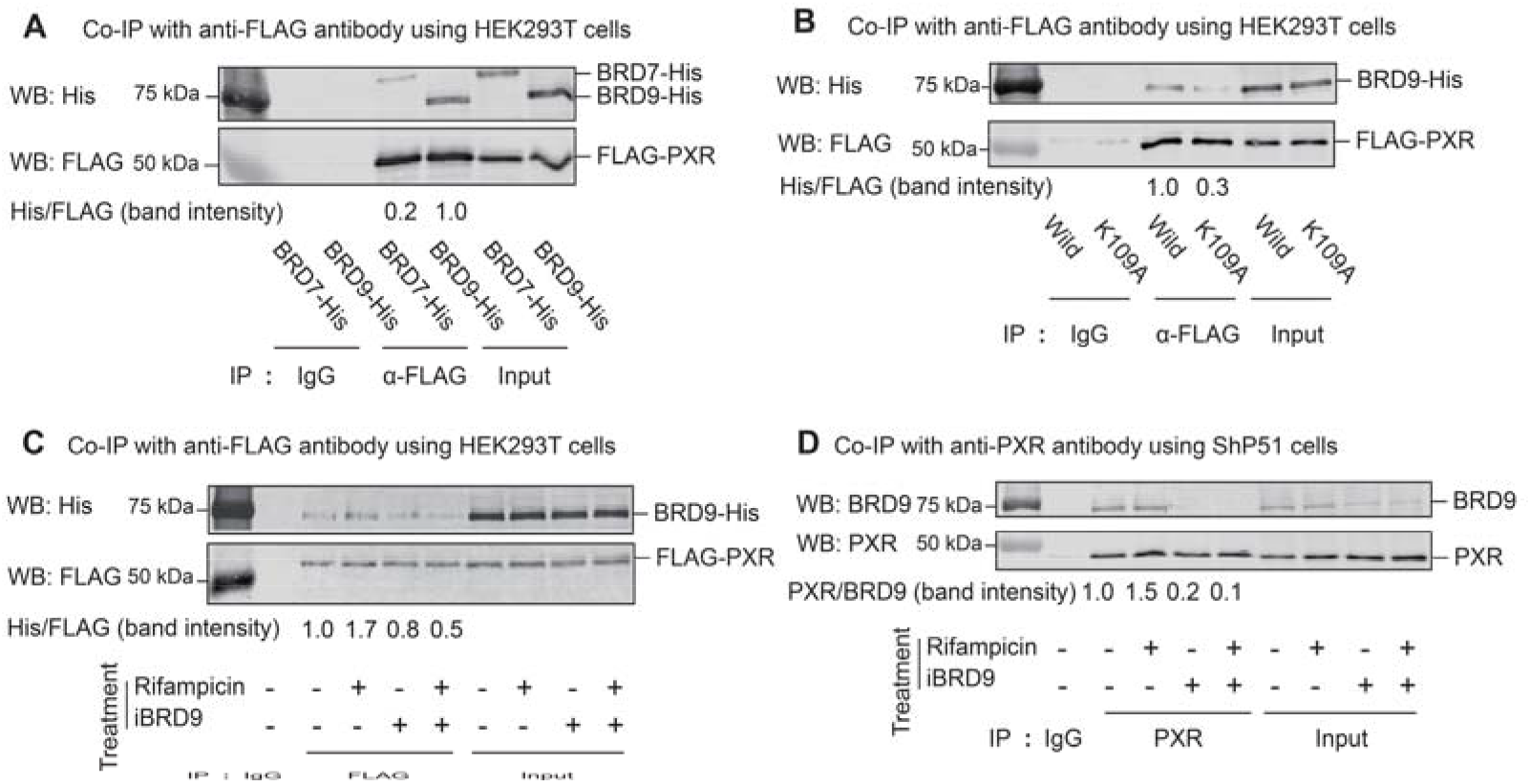
Interaction between PXR and BRD9. HEK293T cells were transfected with FLAG-PXR along with BRD7-His or BRD9-His plasmid (A), transfected with BRD9-His along with FLAG-PXR or FLAG-PXR K109A plasmid (B), or transfected with FLAG-PXR and BRD9-His plasmids followed by treatment with rifampicin along with iBRD9 for 1 hr (C). ShP51 cells were treated with rifampicin along with iBRD9 for 24 hr (D). (A, B, and C) Immunoprecipitation with anti-FLAG antibody was performed using whole HEK293T cell lysates, and FLAG and His tags were detected by Western blotting. (D) Immunoprecipitation with anti-PXR antibody was performed using ShP51 cell nuclear extracts, and BRD9 and PXR proteins were detected by Western blotting. IP: immunoprecipitation, WB: Western blotting. The experiments were repeated two times with similar results.

To investigate whether the interaction between PXR and BRD9 depends on the acetylated K109 of PXR, we constructed FLAG-PXR K109A plasmid in which K109 was substituted with alanine (A). Western blotting using anti-FLAG antibody confirmed that the expression level of FLAG-PXR K109A was almost equal to that of FLAG-PXR (Fig. S1B). Western blotting using anti-acetylated lysine antibody demonstrated that the acetylation level in FLAG-PXR K109A was 0.4-fold that in PXR wild type (Fig. S1B). To investigate whether the substitution of PXR K109A affects the interaction between FLAG-PXR and BRD9-His, co-immunoprecipitation using anti-FLAG antibody was performed. As shown in Fig. 2B, the interaction between FLAG-PXR K109A and BRD9-His was weaker (0.3-fold) than that between FLAG-PXR and BRD9-His, indicating that BRD9 interacts with PXR via acetylated K109. Since similar results were obtained by pull-down assay (Fig. S1C), it was suggested that BRD9 directly interacts with PXR via acetylated K109.

Next, we examined whether the interaction of PXR and BRD9 is modulated by a ligand of PXR and is dependent on the bromodomain of BRD9. FLAG-PXR and BRD9-His-transfected HEK293T cells were treated with rifampicin (a typical PXR ligand) and/or iBRD9 (a selective inhibitor of the bromodomain of BRD9) (26), and co-immunoprecipitation using anti-FLAG antibody was performed. As shown in Fig. 2C, the interaction between FLAG-PXR and BRD9-His was enhanced by rifampicin by 1.7-fold, and the increased interaction was attenuated by iBRD9 to 0.5-fold. The same experiment was performed by using lysates from ShP51 cells, and it was demonstrated that endogenous PXR and BRD9 interact (Fig. 2D). The interaction was increased by 1.5-fold by rifampicin and was decreased by iBRD9 to 0.2-fold (Fig. 2D). As shown in Fig. S1D, rifampicin did not affect the acetylation level of PXR. These results suggested that 1) PXR interacts with BRD9 via the bromodomain of BRD9, 2) the interaction is enhanced by rifampicin, and 3) the enhancement is not due to the increased acetylation level of PXR.

### Effects of knockdown or inhibition of BRD9 on PXR-mediated transactivation

To investigate the role of BRD9 in PXR-mediated transactivation, ShP51 cells were transfected with siBRD7 or siBRD9 followed by treatment with rifampicin or simvastatin. The BRD7 and BRD9 protein levels were significantly (*P* < 0.05) decreased by transfection of siBRD7 and siBRD9, respectively (Fig. S2A). The induction of CYP3A4 mRNA, a downstream gene of PXR, by rifampicin was significantly (*P* < 0.001) attenuated by knockdown of BRD9 but not by knockdown of BRD7 (Fig. 3A). The induction of CYP3A4 mRNA by simvastatin was decreased by approximately half by knockdown of BRD7 and was completely abrogated by knockdown of BRD9. PXR and RXRα protein levels were not changed by treatment with rifampicin or simvastatin and/or knockdown of BRD7 or BRD9 (Fig. 3A). These results suggest that ncBAF containing BRD9 positively regulates PXR-mediated transactivation.

**Figure 3.**
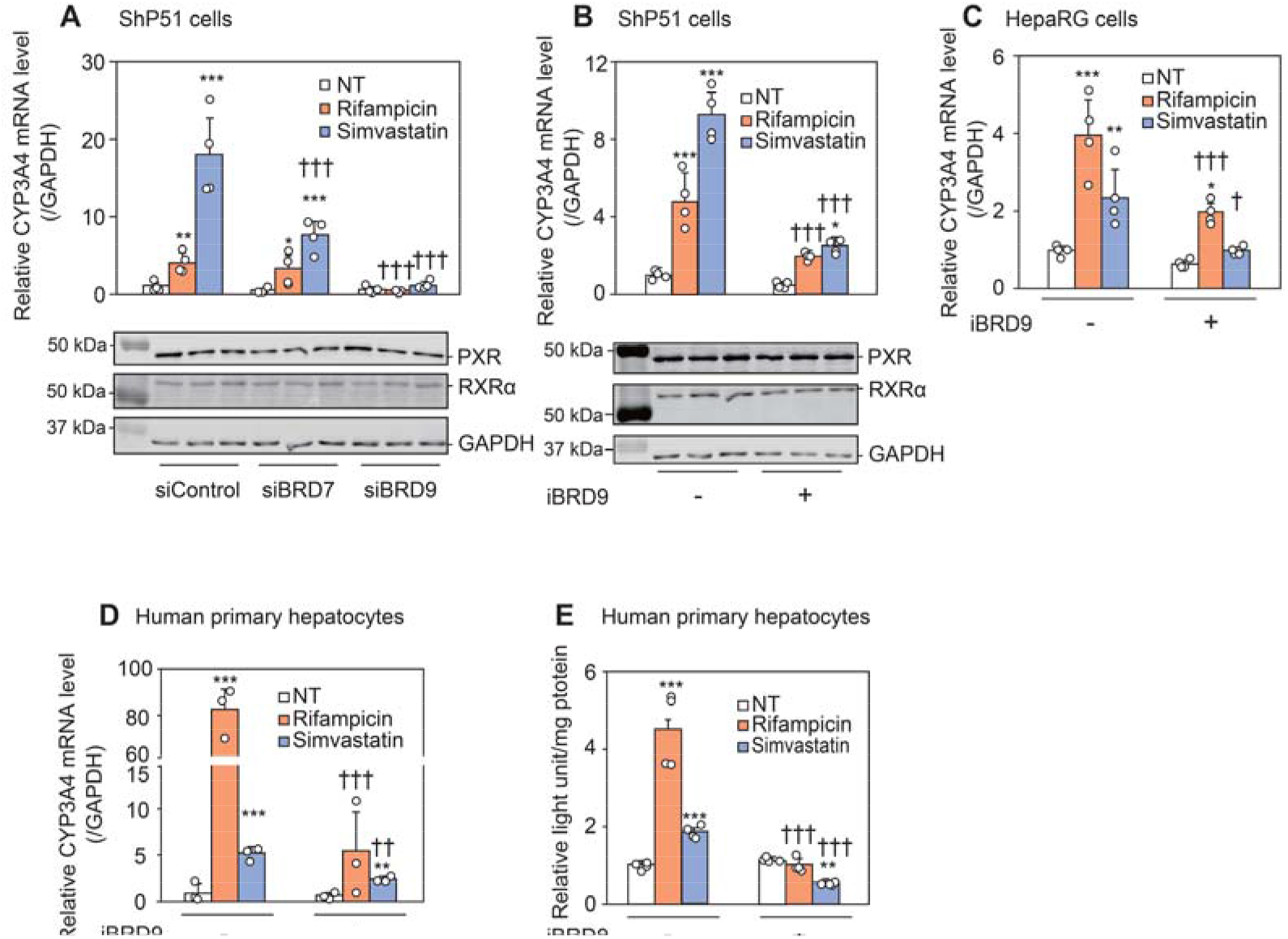
Effects of BRD7 or BRD9 knockdown or BRD9 inhibition on PXR-mediated induction of CYP3A4 expression. (A) ShP51 cells were transfected with siRNA for BRD7 (siBRD7) or BRD9 (siBRD9). After incubation for 24 hr, the cells were treated with 10 μM rifampicin or 10 μM simvastatin. ShP51 cells (B), HepaRG cells (C), and human primary hepatocytes (D and E) were treated with rifampicin or simvastatin along with iBRD9. PXR, RXRα and GAPDH (A and B) protein, CYP3A4 and GAPDH mRNA (A-D), and CYP3A4 enzyme activity (E) were evaluated by Western blotting, real-time RT□PCR, and P450-Glo assay. Each column represents the mean ± SD (n =3-4). n refers to biological repeats. \**P* < 0.05, \*\**P* < 0.01, and \*\*\**P* < 0.001, compared with NT, ^†^*P* < 0.05, ^††^*P* < 0.01, and ^†††^*P* < 0.001, compared with siControl or iBRD9 (−). NT: non-treatment. The experiments were repeated two times with similar results.

Next, we examined whether PXR-mediated induction of CYP3A4 is affected by iBRD9 in ShP51 cells. The induction of CYP3A4 mRNA by rifampicin or simvastatin was significantly (*P* < 0.001) attenuated by iBRD9 treatment in ShP51 cells (Fig. 3B). A question arose regarding whether iBRD9 may affect the translocation of PXR from the cytosol to the nucleus. It has been reported that PXR, in monolayer HepG2 cells or PXR-overexpressing cells, is localized in the nucleus even under exogenous ligand-non-treated condition, while PXR, in 3D-cultured cells, translocates from the cytosol to the nucleus upon ligand treatment (27). To investigate whether iBRD9 affects PXR translocation from the cytosol to the nucleus, 3D culture of ShP51 cells was conducted by using micro-space cell culture plate or hanging drop plate. In both 3D-cultured cells, the induction of CYP3A4 by rifampicin or simvastatin was significantly attenuated by iBRD9 (Fig. S2B), consistent with the results in Fig. 3B. By immunofluorescence staining of PXR in spheroids of ShP51 cells cultured in the hanging drop plate, it was demonstrated that PXR translocated from cytoplasm to nucleus upon rifampicin or simvastatin treatment, and the translocation was not affected by iBRD9 (Fig. S2C). These results suggest that BRD9 regulates PXR-mediated transcriptional activation without affecting the nuclear translocation of PXR. To investigate whether the inhibition of BRD9 also affects the activation of endogenous PXR, the same experiment was performed using HepaRG cells, which relatively retain P450 expression and activity comparable to those in human primary hepatocytes (28). As shown in Fig. 3C, similar results were obtained with ShP51 cells. Finally, to examine whether iBRD9 also affects the induction of CYP3A4 in normal hepatocytes, experiments using primary human hepatocytes were performed. CYP3A4 mRNA level (Fig. 3D) and activity (Fig. 3E) were significantly (*P* < 0.001) increased by rifampicin or simvastatin, and the increases were significantly (*P* < 0.01) attenuated by iBRD9. These results indicate that the inhibition of BRD9 attenuates not only exogenous but also endogenous PXR-mediated transcriptional activation in hepatocarcinoma-derived cells and normal hepatocytes.

We investigated whether the inhibition of BRD9 affects the induction of other PXR downstream genes, CYP2B6, CYP2C9, and UGT1A1 mRNA levels in ShP51 cells and human primary hepatocytes. As shown in Fig. S2D and S2E, although CYP2B6 mRNA level in ShP51 cells was not detected, that in human primary hepatocytes was increased by rifampicin (*P* < 0.001), and the induction was attenuated by iBRD9 (*P* < 0.001). CYP2C9 mRNA levels in ShP51 cells and human primary hepatocytes were increased by rifampicin or simvastatin to varying degrees, and the increases were attenuated by iBRD9. The UGT1A1 mRNA level in ShP51 cells was significantly (*P* < 0.001) increased by rifampicin or simvastatin, and the induction was not attenuated by iBRD9, although the UGT1A1 mRNA level in human primary hepatocytes was not changed by treatment with rifampicin or simvastatin along with iBRD9. Taken together, these results suggested that PXR-mediated transactivation of a certain set of downstream genes requires ncBAF, although PXR-mediated transcriptional activation of UGT1A1 does not require ncBAF.

### BRD9 is required for the binding of PXR to the target region and the change in chromatin structure

To investigate whether the inhibition of BRD9 and activation of PXR affect the binding of PXR to its response elements on a genome-wide scale, sequential salt extraction (SSE) analysis was performed, followed by Western blotting using anti-PXR and BRD9 antibodies. As shown in Fig. 4A, BRD9 in non-treated sample was mainly eluted in the 300 mM NaCl fraction. Rifampicin treatment did not alter the affinity of BRD9 for chromatin but slightly increased the band intensities of BRD9, suggesting that activation of PXR recruits BRD9 to chromatin. The peak of the band density of BRD9 in the iBRD9-treated sample was shifted to the 250 mM NaCl fraction, and the total band intensity of BRD9 was decreased by iBRD9 treatment, suggesting that iBRD9 blocks the binding of BRD9 to chromatin. Similarly, PXR in non-treated sample was mainly eluted in the 300 mM NaCl fraction, and rifampicin treatment did not alter the affinity of PXR for chromatin. Rifampicin treatment increased the total band intensity of PXR, indicating that the activation of PXR promotes the binding of PXR to chromatin. The peak of the band density of PXR in the iBRD9-treated sample was shifted to the 250 mM NaCl fraction, and the increase in total band intensities of PXR by rifampicin was attenuated by iBRD9. These results suggested that ncBAF is required for the binding of PXR to chromatin.

**Figure 4.**
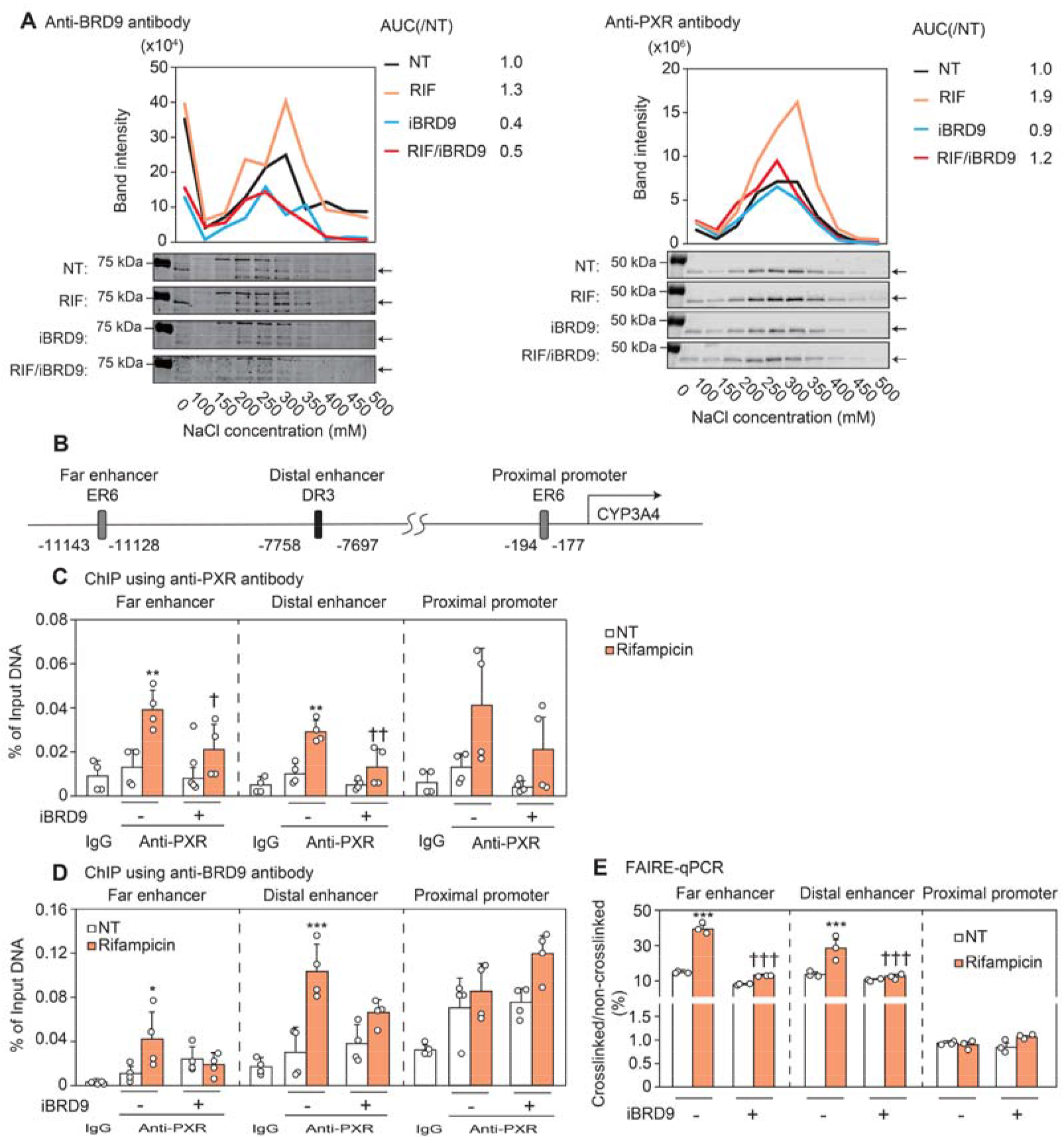
Effects of iBRD9 on PXR binding to chromatin and changes in chromatin structure. (A) ShP51 cells were treated with 20 μM rifampicin along with 20 μM iBRD9 for 24 hr. SSE analysis followed by Western blotting for BRD9 or PXR was performed. The peaks of band intensity are shown in red. (B) Schematic representation of the upstream of *CYP3A4* gene. ShP51 cells were treated with 10 μM rifampicin along with 10 μM iBRD9 for 24 hr. Immunoprecipitation with anti-PXR antibody (C) or anti-BRD9 antibody (D) was performed using chromatin from ShP51 cells. Enrichment of the proximal promoter, distal enhancer, and far enhancer of CYP3A4 was evaluated by real-time PCR. (E) Changes in chromatin structure was evaluated by FAIRE followed by real-time PCR. Each column represents the mean ± SD (n = 3). n refers to biological repeats. \**P* < 0.05, \*\**P* < 0.01, and \*\*\**P* < 0.001, compared with NT, ^††^*P* < 0.01 and ^†††^*P* < 0.001, compared with iBRD9 (−). NT: non-treatment. The experiments were repeated two times with similar results.

PXR is known to bind to everted repeat (ER) 6 in the proximal promoter region (−194/−177), direct repeat (DR) 3 in the distal enhancer region (−7,758/−7,697), and ER6 in the far enhancer region (−11,143/−11,129) of the *CYP3A4* gene (3, 29) to enhance CYP3A4 transcription (Fig. 4B). To investigate whether the inhibition of BRD9 prevents the binding of ligand-activated PXR to the 5’-flanking region of *CYP3A4*, we performed ChIP assay with anti-PXR antibody using rifampicin and/or iBRD9-treated ShP51 cells. As shown in Fig. 4C, the binding of PXR to the far enhancer and the distal enhancer regions were significantly (*P* < 0.01) increased by rifampicin, and the increases were significantly (*P* < 0.05) attenuated by iBRD9, suggesting that ncBAF facilitates the binding of PXR to the 5’-flanking region of *CYP3A4*. To investigate whether the ligand-dependent enhancement of the interaction between PXR and BRD9 (shown in Fig. 2C and D) leads to recruitment of BRD9 to the 5’-flanking region of *CYP3A4*, ChIP assay with anti-BRD9 antibody using rifampicin and/or iBRD9-treated ShP51 cells was performed. The binding of BRD9 to the far and distal enhancer regions were increased by rifampicin, and the increases were attenuated by iBRD9 treatment (Fig. 4D). In the proximal promoter region, the binding of BRD9 was not changed by rifampicin and/or iBRD9 treatment (Fig. 4D). To investigate whether the recruitment of ncBAF in the process of PXR activation leads to changes in chromatin structure, we performed FAIRE assay for the 5’-flanking region of the *CYP3A4* gene. Chromatin structures around the far enhancer and distal enhancer regions were relaxed by rifampicin, and iBRD9 attenuated these structural changes, but that around the proximal promoter region was not changed by rifampicin and/or iBRD9 (Fig. 4E). These results suggest that PXR recruits ncBAF to the far enhancer and distal enhancer regions of *CYP3A4*, resulting in relaxation of chromatin structures.

### iBRD9 attenuates PXR-mediated transcriptional activation in vivo

Next, we investigated by using C57BL/6J mice whether the inhibition of BRD9 affects PXR-mediated transcriptional activation *in vivo*. Because the amino acid identity in the ligand binding domains between mouse Pxr and human PXR is 77%, PCN activates mouse Pxr, while rifampicin does not (30). Accordingly, PCN was administered to mice along with iBRD9. As shown in Fig. 5A, the induction of Cyp3a11 and Cyp3a25 mRNAs, downstream genes of mouse Pxr, by PCN was significantly (*P* < 0.05) attenuated by iBRD9 co-treatment. The induction of Cyp3a protein level (Fig. 5B) and triazolam α- and 4-hydroxylase activities (Fig. 5C) by PCN was significantly (*P* < 0.05) attenuated by iBRD9 co-treatment, suggesting that the inhibition of BRD9 attenuates Pxr-mediated transcriptional activation *in vivo*.

**Figure 5.**
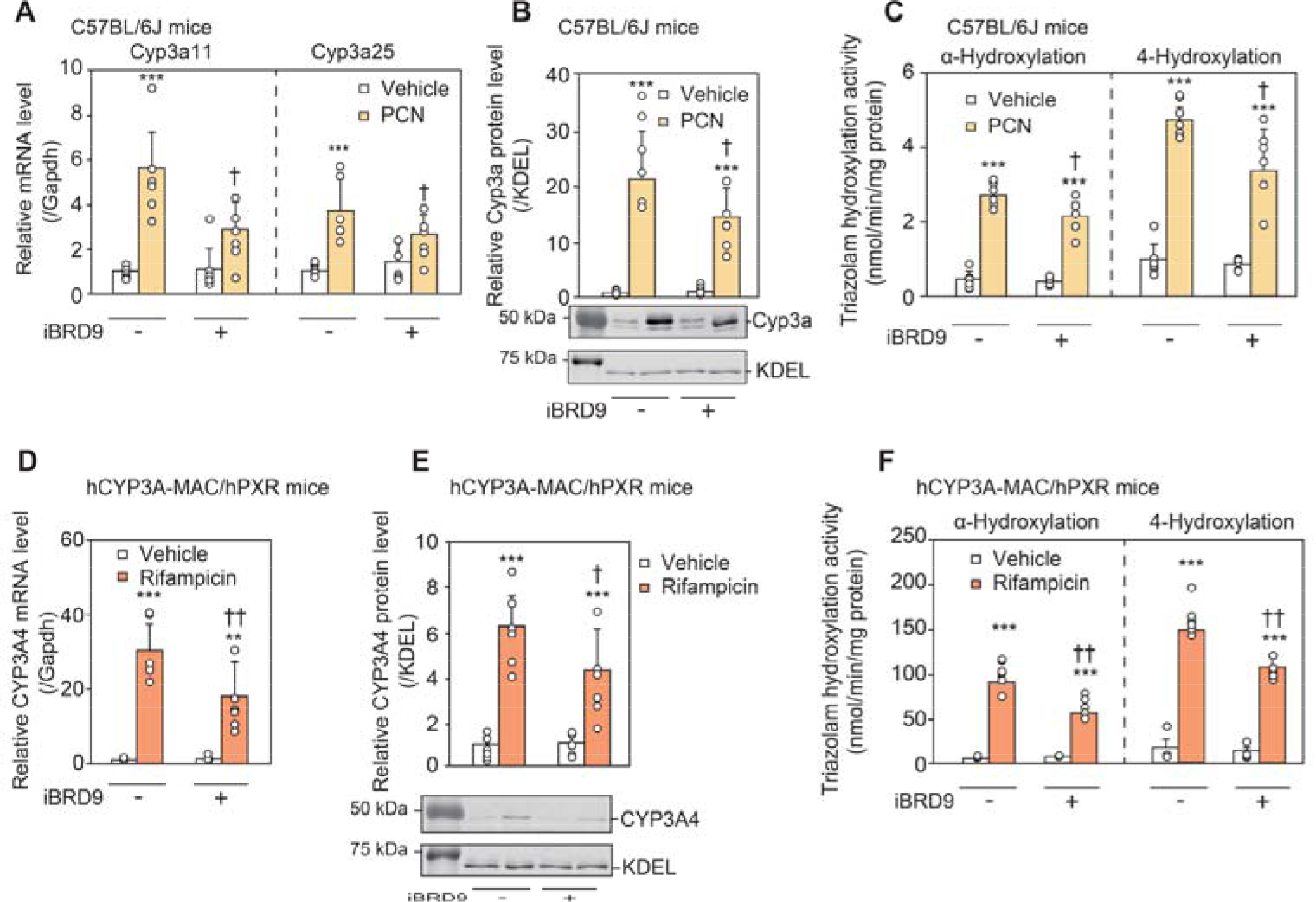
Effects of iBRD9 on the induction of CYP3A in mouse liver. C57BL/6J or hCYP3A-MAC/hPXR male mice (n = 6) were intraperitoneally treated with 50 mg/kg PCN or 10 mg/kg rifampicin for four consecutive days and intraperitoneally treated with 10 mg/kg iBRD9 every other day. Cyp3a11, Cyp3a25, CYP3A4, and Gapdh mRNA (A, D), and Cyp3a, CYP3A4, and KDEL protein (B, E) levels were evaluated by real-time RT□PCR and Western blotting, respectively. (C, F) Triazolam α- and 4-hydroxylase activities were evaluated as marker activity for Cyp3a and CYP3A4. Each column represents the mean ± SD (n = 6). n refers to biological repeats. \*\**P* < 0.01 and \*\*\**P* < 0.001, compared with vehicle, †*P* < 0.05 and ††*P* < 0.01, compared with iBRD9 (−). The experiments were repeated two times with similar results.

Next, hCYP3A-MAC/hPXR mice were used to examine whether iBRD9 attenuates human PXR-mediated transcriptional activation of human CYP3A4 *in vivo*. The mice were treated with rifampicin along with iBRD9. The hepatic CYP3A4 mRNA level (Fig. 5D), protein level (Fig. 5E), and triazolam α- and 4-hydroxylase activities (Fig. 5F) were significantly (*P* < 0.001) increased by rifampicin, and the increases were significantly (*P* < 0.05) attenuated by iBRD9 co-treatment. These results indicate that ncBAF positively regulates mouse and human PXR-mediated transactivation *in vivo*.

The effects of BRD9 inhibition on the induction of other PXR downstream genes, Cyp2b10, Cyp2c29, and Ugt1a1 mRNA levels, were evaluated in C57BL/6J and hCYP3A-MAC/hPXR mice. Cyp2b10 and Cyp2c29 mRNA levels were significantly (*P* < 0.05) increased by PCN and rifampicin in C57BL/6J mice (Fig. S3A) and hCYP3A-MAC/hPXR mice (Fig. S3B), respectively, and the increases tended to be attenuated by iBRD9, although Ugt1a1 mRNA levels in both mice were not changed by PXR ligands. These results suggest that ncBAF regulates mouse and human PXR-mediated transactivation *in vivo* consistent with Fig. S2D and S2E.

### iBRD9 can potentially suppress the risk of simvastatin-induced gluconeogenesis and efavirenz-induced dyslipidemia

It is known that simvastatin induces hyperglycemia (8). Simvastatin-mediated activation of PXR causing the increased hepatic expression of gluconeogenic enzymes, G6Pase and PEPCK1, has been suggested as one of the mechanisms (9). In the present study, we investigated whether the inhibition of BRD9 can suppress the simvastatin-mediated increased expression of gluconeogenic enzymes and glucose production. As shown in Fig. S4A, G6Pase and PEPCK1 mRNA levels in ShP51 cells were significantly (*P* < 0.001) increased by simvastatin, and the inductions were significantly (*P* < 0.001) attenuated by iBRD9. In addition, the increased glucose production by simvastatin was significantly (*P* < 0.01) attenuated by iBRD9 (Fig. S4B), indicating that the inhibition of BRD9 attenuates the gluconeogenesis increase caused by simvastatin. In human primary hepatocytes, the increased G6Pase mRNA level induced by simvastatin was not attenuated but rather enhanced by iBRD9 (Fig. S4C). PEPCK1 mRNA level was significantly (*P* < 0.01) increased by simvastatin, and the induction was decreased by iBRD9. These results suggest that inhibition of BRD9 attenuates PXR-mediated hyperglycemia, although ncBAF appears to be strongly affected by factors other than PXR in the case of G6Pase in human primary hepatocytes.

Efavirenz has been reported to increase the risks of hypercholesterolemia and hepatic steatosis in the clinic (31). A recent report has shown that efavirenz activates human PXR and mouse Pxr, resulting in the induction of expressions of SQLE, a cholesterol synthase, and CD36, a hepatic fatty acid uptake transporter (11). To investigate whether efavirenz-induced dyslipidemia and hepatic steatosis can be suppressed by the inhibition of BRD9, C57BL/6J mice were treated with efavirenz along with iBRD9. Unexpectedly, 7 out of 12 mice that received efavirenz died within 7 days (Fig. 6A), even though the dose administration route and schedule of efavirenz were the same as those reported previously (11). Two out of eight mice that received efavirenz/iBRD9 died, and their survival rate was higher than that of efavirenz-treated mice. In subsequent experiments, the effects of BRD9 inhibition on the efavirenz-mediated transactivation of PXR were examined in 6 vehicle-treated, 5 efavirenz-treated, and 6 efavirenz/iBRD9-treated mice. As shown in Fig. 6B, the Cyp3a11 mRNA level was significantly (*P* < 0.001) increased by efavirenz, and the increase was significantly (*P* < 0.05) attenuated by iBRD9, suggesting that efavirenz-mediated activation of Pxr can be suppressed by iBRD9. Sqle and Cd36 mRNA levels were significantly (*P* < 0.05) increased by efavirenz, and the levels were attenuated by iBRD9. Interestingly, the liver weight was increased by efavirenz, and the increase tended to be attenuated by iBRD9 (Fig. 6C). It has been reported that efavirenz increases plasma cholesterol level and decreases plasma triglyceride level (11), and such trends were observed in the present study (Fig. 6D). Consistent with a previous report, hepatic cholesterol and triglyceride levels were increased by efavirenz, and the changes were attenuated by iBRD9 (Fig. 6D). In addition, hepatic lipid accumulation evaluated by oil red O staining was induced by efavirenz, and the increase was attenuated by iBRD9 (Fig. 6E). These results suggest that the inhibition of BRD9 can suppress efavirenz-induced hepatic steatosis by PXR activation.

**Figure 6.**
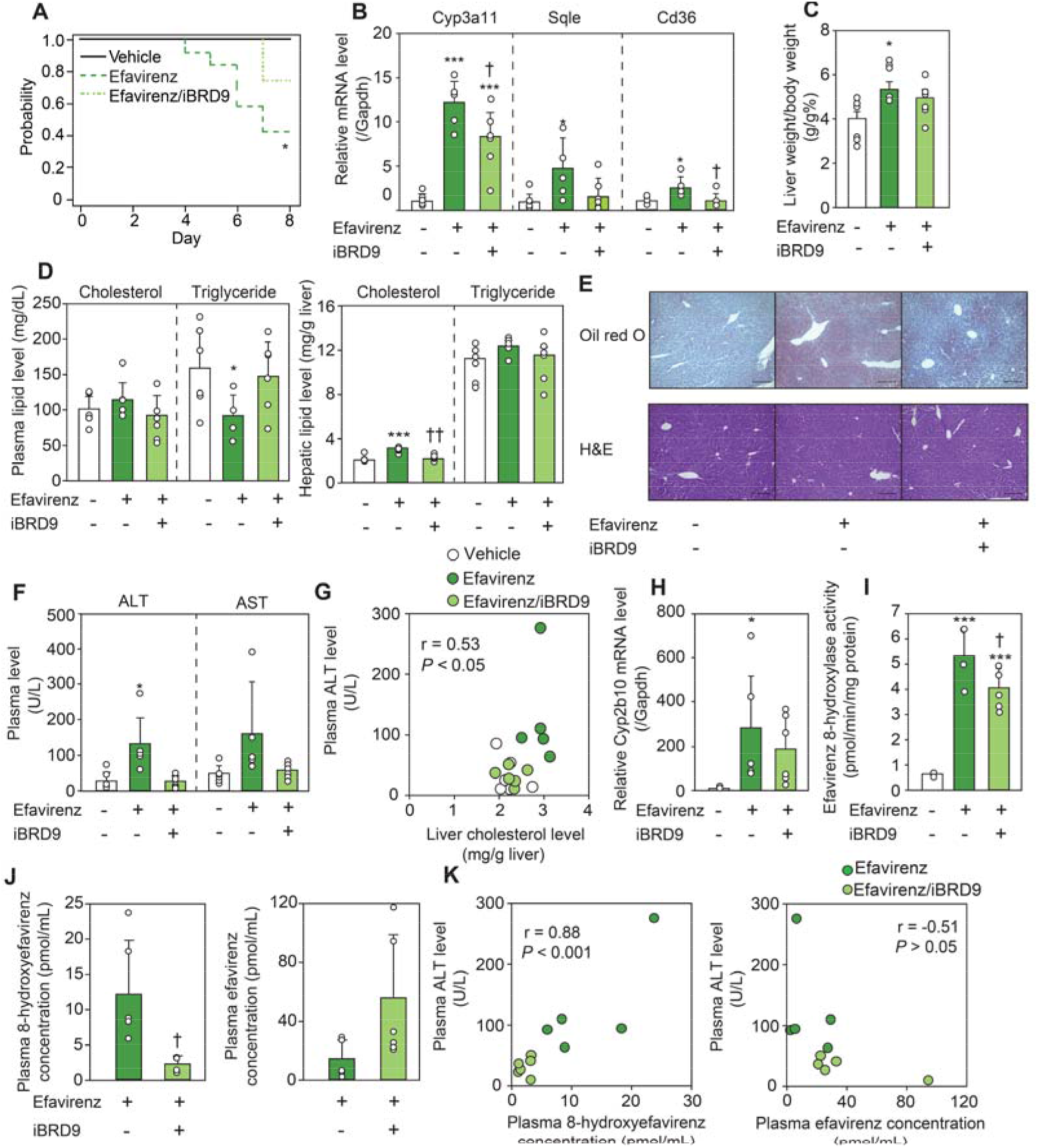
Effects of iBRD9 treatment on efavirenz-mediated hepatic lipid accumulation and hepatotoxicity *in vivo*. C57BL/6J male mice fed a low-fat diet for 1 week were orally treated with 20 mg/kg efavirenz and intraperitoneally treated with 20 mg/kg iBRD9 daily for 1 week. (A) Survival rate of mice administered with efavirenz and/or iBRD9. (B) Cyp3a11, Sqle and Cd36 mRNA levels were evaluated by real-time RT□PCR. (C) The ratio of liver weight to body weight. Plasma and hepatic cholesterol and triglyceride (D) and plasma AST and ALT (F) levels were evaluated. (E) Liver specimens were subjected to H&E staining and oil red O staining. Correlation of plasma ALT level with hepatic cholesterol level (G), plasma efavirenz level or plasma 8-hydroxy efavirenz level (K). (H) Cyp2b10 mRNA level was evaluated by real-time RT□PCR. (I) Efavirenz 8-hydroxylase activity was evaluated as a marker activity for Cyp2b10. (J) Plasma 8-hydroxyefavirenz and efavirenz levels were evaluated. Each column represents the mean ± SD (n = 5–6). n refers to biological repeats. \**P* < 0.05 and \*\*\**P* < 0.001, compared with vehicle, ^†^*P* < 0.05 and ^††^*P* < 0.01, compared with efavirenz administration. The experiments were repeated two times with similar results.

Next, to obtain an answer to the question of why iBRD9 can reduce the mortality caused by efavirenz (Fig. 6A), the effects of iBRD9 on efavirenz-induced hepatotoxicity were examined. Plasma ALT level was increased by efavirenz, and the increase was restored by iBRD9. Plasma AST level showed a similar tendency (Fig. 6F). These results suggest that the mitigation of efavirenz-induced hepatotoxicity by iBRD9 might be one of the reasons for the decreased mortality. Considering the possibility that hepatotoxicity was caused by lipid accumulation, the relationship between hepatic cholesterol content and ALT level was examined. A significant correlation was observed between them (*P* < 0.05), but the correlation was moderate (r = 0.53) (Fig. 6G), suggesting the involvement of factor(s) other than lipid accumulation in efavirenz-induced hepatotoxicity. It has been reported that 8-hydroxyefavirenz, a main metabolite of efavirenz, is more toxic than efavirenz to primary human hepatocytes (32). The 8-hydroxylation of efavirenz is mainly catalyzed by CYP2B6 (33), of which the mouse ortholog is Cyp2b10. Hepatic Cyp2b10 mRNA level (Fig. 6H) and efavirenz 8-hydroxylase activity (Fig. 6I) were significantly (*P* < 0.05) increased by efavirenz, and the increases were attenuated by iBRD9. As shown in Fig. 6J, the plasma 8-hydroxyefavirenz concentration in efavirenz/iBRD9-treated mice was significantly (*P* < 0.05) lower than that in efavirenz-treated mice, whereas the plasma efavirenz concentration in efavirenz/iBRD9-treated mice tended to be higher than that in efavirenz-treated mice. The ALT level was significantly (*P* < 0.001) correlated (r = 0.88) with the plasma 8-hydroxyefavirenz concentration but not with the efavirenz concentration (Fig. 6K), suggesting that the extent of efavirenz 8-hydroxylation is a major determinant of efavirenz-induced hepatotoxicity. Collectively, we clarified that the adverse events caused by efavirenz can be attenuated by the inhibition of BRD9.

## Discussion

Chromatin remodeling is essential for the transcription, replication, and recombination of DNA (20). Dynamic structural changes in chromatin are caused by chromatin remodelers such as the SWI/SNF complex (34). In the 2000s, BAF60a and BAF57, components of the SWI/SNF complex, were reported to interact with GR, leading to up-regulation of GR-mediated transcriptional activation (35). These components are commonly included in cBAF, PBAF, and ncBAF. The roles of the other sub-complex-specific components in nuclear receptor-mediated transcription were not studied in the early 2000s. Recently, it has become clear that PBAF and ncBAF differentially regulate VDR-mediated transcriptional regulation through recognition by BRD7 and BRD9, respectively (36). More recently, it has been reported that BRD9 positively and negatively regulates AR- and GR-mediated transcriptional activation, respectively (37–38). The present study sought to clarify the roles of the SWI/SNF complex in transcriptional activation by PXR, which belongs to the nuclear receptor superfamily along with VDR, AR, and GR and plays an important role in the regulation of drug metabolism.

By the co-immunoprecipitation using anti-PXR antibody and glycerol gradient sedimentation, the interaction of PXR with ncBAF via BRD9 was demonstrated (Figs. 1 and 2). Contrary to the results, BRD9 was not detected in immunoprecipitants using anti-PXR antibody by mass spectrometric analysis (Fig. 1A). It is known that LC□MS-based proteomics may fail to identify proteins if the conditions are inappropriate (39). It has also been reported that the reproducibility of protein detection between laboratories was low in some cases, even though the same sample was measured (40). The fact that BRD9 was not detected in the immunoprecipitant is probably because the proteomic conditions used were not suitable for BRD9.

Co-immunoprecipitation analysis indicated dependence of the interaction between PXR and BRD9 on the bromodomain of BRD9 and the acetylated K109 residue of PXR (Fig. 2B). It has been reported that the acetylation level of the lysine residue of mouse Pxr is increased by feeding of high-fat diet (41). The elevation of the acetylated-lysine level would enhance the interaction between PXR and BRD9, potentially leading to enhancement of PXR-mediated transactivation of drug-metabolizing enzymes, fatty acid synthases such as fatty acid elongase and stearoyl-CoA desaturase-1, and cholesterol synthases such as SQLE in the liver (42). Therefore, this post-translational modification of PXR may affect PXR-mediated drug□drug interaction and obesity (43). The present study demonstrated that the acetylation level of PXR was not changed by rifampicin (Fig. S1D). The K109 sits in the hinge domain, which has been simulated to be flexible (25). The increased interaction between PXR and BRD9 induced by rifampicin may be due to ligand-mediated conformational changes in the PXR protein. A computational approach would be merited to verify the assumption.

As with the case of CYP3A4, the inductions of CYP2B6, CYP2C9, G6Pase, and PEPCK1 mRNA levels by ligands in ShP51 cells and human primary hepatocytes were significantly attenuated by knockdown of BRD9 or co-treatment with iBRD9 (Fig. 3 and Fig. S2). On the other hand, the PXR-mediated induction of UGT1A1 in ShP51 cells was not changed by iBRD9. These results indicate that ncBAF does not enhance all PXR-mediated transactivation, and other regulatory mechanisms may be involved in the regulation of UGT1A1 transcription. A recent study revealed that ncBAF uniquely localizes to CCCTC-binding factor (CTCF) binding sites and promoter regions (21). According to the ChIP atlas database (http://chip-atlas.org), CTCF binds to the flanking regions within 20 kb of each *CYP2B6*, *CYP2C9*, *G6Pase*, *PEPCK1*, or *CYP3A4* gene but not to the flanking region of the *UGT1A1* gene (Fig. S5). This fact may imply that ncBAF is responsible for the enhancement of transactivation of PXR-targeted genes accompanied by the CTCF binding region. Further studies are needed to clarify the possibility that CTCF may affect the function of ncBAF interaction with nuclear receptors.

PXR transcriptionally activates the *CYP3A4* gene by binding to its responsive sequences in the 5’-upstream region (13–15). FAIRE assay and ChIP assay using anti-PXR or anti-BRD9 antibody demonstrated that ncBAF relaxes the chromatin structure around the enhancer regions of *CYP3A4*, facilitating PXR binding to response elements (Fig. 4). These results suggest that PXR recruits ncBAF to the enhancer regions of *CYP3A4*, resulting in chromatin decondensation. FAIRE-seq data compiled in the UCSC genome browser (https://genome.ucsc.edu/) reveal that the chromatin structure in the proximal promoter region of *CYP3A4* is relatively open in comparison with the enhancer regions. This means that structural changes in chromatin in the vicinity of this region would not be required for transactivation. This might be the reason why the binding of BRD9 to the promoter region was not changed by iBRD9 or rifampicin treatment (Fig. 4D).

Inhibitors of PXR are expected to reduce PXR-mediated adverse events. SPA70 has been reported as a PXR inhibitor that binds to the ligand recognition site of PXR to prevent ligand binding, resulting in repression of the induction of downstream genes of PXR by rifampicin or SR12813 in human primary hepatocytes (44). However, this inhibitor stabilizes the PXR protein, resulting in an approximately 2-fold increase in the basal expression level of CYP3A4 (44). In the case of iBRD9, it did not change the basal expression levels of drug-metabolizing enzymes, although it suppressed ligand-activated PXR function (Fig. 3, Fig. 5, Fig. S2, and Fig. S3), probably because iBRD9 inhibits the interaction between PXR and BRD9 but does not directly bind to the ligand recognition site. In addition, iBRD9 attenuated simvastatin-induced gluconeogenesis and efavirenz-induced hepatic steatosis without affecting the basal expression levels of gluconeogenesis enzymes and lipid synthesis enzymes (Fig. 6 and Fig. S4). Therefore, iBRD9 would be useful to reduce the risk of PXR-mediated adverse events such as drug□drug interaction, hyperglycemia, and dyslipidemia.

Accumulating evidence indicates that BRD9 plays an oncogenic role in multiple cancer types by regulating tumor cell growth (45). Therefore, BRD9 is an attractive target for cancer therapy. In addition, the biological significance of BRD9 in β cell function and inflammation has been revealed (36, 38). The present study demonstrated the roles of BRD9 in the regulation of normal liver function, including drug, glucose, and lipid metabolism. Although iBRD9 did not completely attenuate the induction of Cyp3a11 (Fig. 6B), it completely abolished efavirenz-induced hepatic lipid accumulation (Fig. 6B, D and E), suggesting that some mechanisms other than PXR-mediated transcription may be involved in the regulation of lipid metabolism by BRD9. More comprehensive research is needed to understand the functional significance of BRD9 in hepatic energy metabolism.

In conclusion, we found that ncBAF is a critical factor regulating transactivation by PXR. This is the first study demonstrating a chromatin remodeler responsible for PXR-mediated transcriptional regulation. Inhibition of BRD9 has the potential to reduce the clinical risk of PXR-mediated adverse events.

## Materials and Methods

### Chemicals and reagents

Rifampicin, triazolam, and α-hydroxytriazolam were purchased from FUJIFILM Wako Pure Chemical (Osaka, Japan). 4-hydroxytriazolam was purchased from Biomol GmbH (Hamburg, Germany). Simvastatin and iBRD9 were purchased from Toronto Research Chemicals (Toronto, Canada) and Cayman Chemical (Ann Arbor, MI), respectively. Pregnenolone 16α-carbonitrile (PCN) and dimethyl pimelimidate were purchased from Medchem Express (Monmouth Junction, NJ) and Tokyo Chemical Industry (Tokyo, Japan), respectively. Efavirenz and 8-hydroxyefavirenz were purchased from Tokyo Chemical Industry and Advanced ChemBlocks (Hayward, CA), respectively. Glucose-6-phosphate (G6P), glucose-6-phosphate dehydrogenase (G6PDH), and β-nicotinamide adenine dinucleotide phosphate oxidized form (NADP^+^) were purchased from Oriental Yeast (Tokyo, Japan). Lipofectamine RNAiMAX, Lipofectamine 3000, Silencer Select siRNA for human BRD7 (s26525) (siBRD7), human BRD9 (s35295) (siBRD9), or negative control #1 (siControl), and Dynabeads protein G were purchased from Thermo Fisher Scientific (Waltham, MA). pTargeT mammalian expression vector and P450-Glo CYP3A4 Assay (Luciferin IPA) were obtained from Promega (Madison, WI). RNAiso and random hexamers were from Takara (Shiga, Japan). ReverTra Ace was purchased from Toyobo (Osaka, Japan). Luna Universal qPCR Master Mix was purchased from New England Biolabs (Ipswich, MA). All primers were commercially synthesized at Eurofins Genomics (Tokyo, Japan). Mouse anti-human PXR (sc-48340), RXRα (sc-515929), CYP3A4 (sc-53850), BRD7 (sc-376180), and Ac-lysine (sc-32268) monoclonal antibodies and the goat anti-mouse Cyp3a (sc-30621) polyclonal antibody were obtained from Santa Cruz Biotechnology (Santa Cruz, CA). Rabbit anti-human BRD9 polyclonal antibody (A303-781A), rabbit anti-ARID1A (GTX129433) antibody, and rabbit anti-SMARCD1 (BAF60A) antibody (10998-2-AP) were obtained from Bethyl Laboratories (Montgomery, AL), Genetex (Irvine, CA), and Proteintech (Rosemont, IL), respectively. Rabbit anti-human GAPDH polyclonal antibody (NB100-56875) and mouse anti-KDEL monoclonal antibody (NBP1-97469) were purchased from Novus Biologicals (Centennial, CO). Mouse anti-FLAG M2 monoclonal antibody (F1804) and mouse anti-His-tag monoclonal antibody (D291-3S) were purchased from Sigma□Aldrich (St. Louis, MO) and Medical and Biological Laboratories (Nagoya, Japan), respectively. IRDye 680 goat anti-rabbit IgG and goat anti-mouse IgG antibodies were purchased from LI-COR Biosciences (Lincoln, NE). All other chemicals and solvents were of the highest grade commercially available.

### Cell cultures

ShP51, a human hepatocellular carcinoma-derived HepG2 cell line stably expressing human PXR (46), was kindly provided by Dr. Negishi (NIH/NIEHS). The cells were cultured in Dulbecco’s modified Eagle’s medium (DMEM, Nissui Pharmaceutical, Tokyo, Japan) containing 10% fetal bovine serum (FBS) and non-essential amino acids (NEAA, Thermo Fisher Scientific). Human hepatocellular carcinoma-derived HepaRG cells, purchased from KAC (Kyoto, Japan), were cultured in William’s E medium supplemented with 10% FBS (Invitrogen), 100 U/mL penicillin, 100 μg/mL streptomycin, 5 μg/mL insulin, 2 mM glutamine, and 50 μM hydrocortisone hemisuccinate. After culturing for 2 weeks, the medium was replaced with the same medium supplemented with 2% DMSO, and the cells were further cultured for 2 weeks to differentiate. The medium was exchanged every 2 or 3 days. Human embryonic kidney-derived HEK293T cells, purchased from American Type Culture Collection (Manassas, VA), were cultured in DMEM supplemented with 10% FBS and 3.5 g/L glucose. Human primary hepatocytes (lot No. HC5-2) obtained from Sekisui Xenotech (Kansas City, KS) were cultured in OptiCULTURE Hepatocyte Culture Media (Sekisui Xenotech). These cells were cultured at 37 °C under an atmosphere of 5% CO_2_ and 95% air.

### Construction and transfection of expression plasmids

FLAG-tagged PXR (FLAG-PXR) plasmid was kindly provided by Dr. Negishi (NIH/NIEHS) (47). FLAG-tagged PXR K109A plasmid was generated by substituting the lysine residue at 109 with alanine by site directed mutagenesis. His-tagged BRD7 (BRD7-His) and BRD9 (BRD9-His) plasmids were constructed by subcloning the BRD7 or BRD9 cDNA with His-tag sequence at their *C*-terminus into a pTarget vector.

HEK293T cells were seeded into 6-well plate. After 24 hr of incubation, the cells were transfected with FLAG-PXR plasmid and BRD7-His or BRD9-His plasmids using Lipofectamine 3000. After 48 hr of incubation, the cells were collected and lysed using ice-cold lysis buffer [20 mM HEPES (pH 8.0), 0.2 mM EDTA, 0.3 M NaCl, 0.5% NP-40, and 15% glycerol].

### Transfection with siRNA and treatment with chemicals followed by preparation of cell homogenates, nuclear extracts, and total RNA

ShP51 cells were seeded and transfected with 5 nM siBRD7 or siBRD9 using Lipofectamine RNAiMAX. After incubation for 24 hr, the ShP51 cells were treated with 10□μM rifampicin or 10□μM simvastatin for 24 hr. In the experiment without transfection with siRNA, ShP51 cells (in monolayer or 3D culture), HepaRG cells, or human primary hepatocytes were treated with 10 μM rifampicin or 10 μM simvastatin along with 10 μM iBRD9 for 24 or 48 hr. The cells were suspended in a small amount of TGE buffer [10 mM Tris-HCl, 20% glycerol, and 1 mM EDTA (pH 7.4)] and disrupted by three freeze-thaw cycles. Nuclear extracts were prepared following a previous report (48) and were resuspended in lysis buffer for subsequent experiments. Protein concentrations were determined according to the Bradford method (49). Total RNA was prepared using RNAiso according to the manufacturers’ protocols.

### Co-immunoprecipitation

Anti-FLAG or PXR antibodies were crosslinked with Dynabeads protein G using dimethyl pimelimidate. The lysates of HEK293T cells or nuclear extracts of ShP51 cells were incubated with the antibody-Dynabeads protein G complex for 2 hr. After washing three times using wash buffer [PBS containing 2 mM EDTA and 0.1% Triton X-100], the beads were incubated with digestion buffer [50 mM triethylammonium bicarbonate (pH 8.5) and 100 μg/mL trypsin], and the supernatant was collected for proteomic analysis or mixed with SDS□PAGE sample buffer [0.2 M Tris-HCl (pH 6.8), 0.2 M SDS, 0.3 mM bromophenol blue, 30% glycerol, and 15% 2-mercaptoethanol] for Western blotting.

### Protein identification using nanoliquid chromatography□mass spectrometry

Protein identification was performed at the Institute for Gene Research (Kanazawa University, Kanazawa, Japan). Briefly, the immunoprecipitant described above was subjected to LC□MS/MS analysis using an Orbitrap QE plus MS (Thermo Fisher Scientific) coupled to an EASY-nLC system (Thermo Fisher Scientific). A trap column, Acclaim PepMap 100 C18 LC column (3 μm, 75 μm × 20 mm, Thermo Scientific P/N16496) and analytical column, NANO-HPLC C18 capillary column (3 μm, 75 μm × 150 mm, Nikkyo Technos, Tokyo, Japan), were used. The columns were equilibrated with 0.1% formic acid, and the sample was eluted with a linear acetonitrile gradient (0–35%) in 0.1% formic acid at flow rate of 300 nL/min. The ion spray voltage was set at 2 kV (ion transfer tube temperature: 275 °C). The peptide ions were detected under the data-dependent acquisition mode using the installed Xcalibur software (Thermo Fisher Scientific). Full-scan mass spectra were acquired in the range of 375-1,500 m/z with resolution of 70,000. The most intense precursor ions were selected for collision-induced fragmentation in the linear ion trap.

### Glycerol gradient sedimentation analysis

Linear 10–30% glycerol gradients were prepared in 11□mm□×□34□mm polyallomer centrifuge tubes (Beckman Coulter, Brea, CA) with HEM buffer [25 mM HEPES (pH 7.5), 0.1 mM EDTA, and 25 mM MgCl_2_]. The nuclear extracts (3□mg protein) resuspended in 150□μL of HEM buffer were overlaid on the top of the tube contents, and the samples were centrifuged with a TLS-55 rotor (Beckman Coulter) at 55,000□rpm at 4 °C for 8□hr. Sixteen 0.1-mL fractions were sequentially collected from the top and subjected to Western blotting.

### SDS□PAGE and Western blotting

The cell homogenates, immunoprecipitants, and MLM were separated by 6% (ARID1A), 7.5% (Ac-Lys, CYP3A4, Cyp3a, RXRα, BRD7, BRD9, SMARCD1, and KDEL) or 10% (GAPDH) SDS□PAGE and transferred to an Immobilon-P transfer membrane (Millipore, Billerica, MA). For PXR, the cell homogenates were separated by 7.5% SDS□PAGE and transferred to a Protran nitrocellulose membrane (Whatman GmbH, Dassel, Germany). The membranes were probed with the primary antibody and then with the fluorescent dye-conjugated secondary antibody. The bands were quantified by using an Odyssey Infrared Imaging system (LI-COR Biosciences). The Cyp3a and CYP3A4 protein levels in C57BL/6J and hCYP3A-MAC/hPXR mice, respectively, were normalized to the KDEL protein level as described in a previous report (50).

### Real-time RT-PCR

The cDNA was synthesized from total RNA using ReverTra Ace according to the manufacturers’ protocols. A 1-μL portion of the reverse-transcribed mixture was added to PCR mixture containing 5□pmol of each primer and 10□μL of Luna Universal qPCR mix in a final volume of 20□μL. Primer sequences for human CYP3A4, CYP2B6, CYP2C9, UGT1A1, G6Pase, PEPCK1, GAPDH as well as mouse Cyp3a11, Cyp3a25, Cyp2b10, Cyp2C29, Ugt1a1, Cd36, Sqle, and Gapdh are shown in Table S1. PCR conditions were as follows: After an initial denaturation at 95°C for 60 s, amplification was performed by denaturation at 95°C for 15 s, then annealing and extension with the conditions as shown in Table S2 for 40 cycles using Mx3000P (Stratagene). Each mRNA level was normalized to the GAPDH mRNA level.

### Measurement of CYP3A4 activity *in cellulo*

*In cellulo* CYP3A4 activity in human primary hepatocytes was measured as follows. Human primary hepatocytes were seeded at 4 × 10^4^ cells/well into 96-well plate. After 3 hr incubation, the cells were treated with 10□μM rifampicin or 10□μM simvastatin along with 10□μM iBRD9. After 48 hr, CYP3A4 activity was measured using the P450-Glo CYP3A4 Assay (Luciferin IPA).

### Sequential salt extraction analysis

SSE analysis was performed following previous report (51) with some modifications. Briefly, ShP51 cells were lysed with modified buffer A [60 mM Tris-HCl (pH 8.0), 60 mM KCl, 1 mM EDTA, 0.3 M sucrose, 0.5%, and 1 mM DTT] and were incubated at 4 °C for 10 min. After centrifugation, the nuclei were incubated in 200 μL of extraction buffer 0 [50 mM Tris-HCl (pH 8.0), 1% NP-40, 0.5% sodium deoxycholate] for 3 min and centrifuged at 6,500 × g for 5 min, and the supernatant was collected as the “0 mM fraction.” The pellet was resuspended in 200 μL of extraction buffer 100 [50 mM Tris-HCl (pH 8.0), 1% NP-40, 0.5% sodium deoxycholate, and 100 mM NaCl] and processed in the same manner to yield the “100 mM fraction”. Serial extraction was implemented with extraction buffers containing 150, 200, 250, 300, 350, 400, 450, and 500 mM NaCl. Twenty-microliter aliquots from each fraction were mixed with SDS□PAGE sample buffer for Western blotting.

### Chromatin immunoprecipitation (ChIP)

ShP51 cells were cross-linked with 1% formaldehyde for 30 min. Chromatin immunoprecipitation assay was performed following a previous report (52). The final DNA samples were analyzed by real-time PCR. Primer sequences and PCR conditions for the proximal promoter, distal enhancer, and far enhancer of *CYP3A4* are shown in Tables S1 and S2.

### Formaldehyde-assisted isolation of regulatory elements assay

ShP51 cells were cross-linked with 1% formaldehyde. Formaldehyde-assisted isolation of regulatory elements was conducted according to a previous report (53). The purified DNA was subjected to real-time PCR using primers designed for the proximal promoter, the distal enhancer, and the far enhancer of *CYP3A4* as described above.

### Administration of PXR ligands and iBRD9 to wild-type and hCYP3A-MAC/hPXR mice

The hCYP3A-MAC/hPXR mice genetically engineered to express human CYP3A4, CYP3A5, CYP3A7, CYP3A43, and PXR were established at Tottori University (54). The hCYP3A-MAC/hPXR mice and control C57BL/6J mice were housed in an institutional animal facility under the controlled environment (temperature 25□±□1 °C and 12 hr light/dark cycle) with access to food and water *ad libitum*. The animal study was approved by the Institutional Animal Care and Use Committees of Kanazawa University and Tottori University and was conducted in accordance with the National Institutes of Health Guide for Animal Welfare of Japan.

The mice (9–11 weeks old, male) were intraperitoneally treated with 50 mg/kg PCN or 10 mg/kg rifampicin for four consecutive days and intraperitoneally treated with 10 mg/kg iBRD9 every other day. Sixteen hours after the final treatment, liver pieces were collected. C57BL/6J mice fed a low-fat diet (Oriental Yeast) for 1 week were orally treated with 20 mg/kg efavirenz and intraperitoneally treated with 20 mg/kg iBRD9 daily for one week. The mice were sacrificed 15 hr after the final treatment, and blood was collected from the inferior vena cava. Mouse liver microsomes (MLMs) were prepared according to a method reported previously (55). Plasma cholesterol, triglyceride, alanine aminotransferase (ALT), and aspartate aminotransferase (AST) levels were measured by DRI-CHEM (Fujifilm Wako Pure Chemical). Hepatic lipid was extracted by Bligh-Dyer method (56), and hepatic triglyceride and cholesterol levels were measured using Lab assay kit (Fujifilm Wako Pure Chemical). Liver sections from mice were fixed in 10% buffered formalin and embedded in paraffin. Then, the sections were sliced, and hematoxylin and eosin (H&E) staining and oil red O staining were performed by the Applied Medical Research Laboratory (Osaka, Japan).

### Measurement of CYP3A4 activity *in vitro*

Triazolam α- and 4-hydroxylase activities were measured as marker activities for mouse Cyp3a and human CYP3A (57). An incubation mixture (final volume of 0.2 mL) contained 100 mM potassium phosphate buffer (pH 7.4), 0.3 mg/mL MLM, and 50 μM triazolam dissolved in methanol (1% v/v final concentration). In a preliminary study, it was confirmed that the triazolam α-hydroxylase activity was linear with respect to protein concentration (≤ 0.5 mg/mL) and incubation time (≤ 60 min). The reaction was initiated by the addition of a NADPH-generating system (0.5 mM NADP^+^, 5 mM G6P, 5 mM MgCl_2_, and 1 U/mL G6PDH) after preincubation at 37°C for 2 min. After a 30-min incubation, the reactions were terminated by the addition of 200 μL of ice-cold acetonitrile. After removal of the protein by centrifugation at 20,380 g for 5 min, a 150 μL portion of the supernatant was subjected to HPLC. HPLC analysis was performed using an L-2130 pump (Hitachi, Tokyo, Japan), an L-2200 autosampler (Hitachi), an L-2400 UV detector (Hitachi), and a D-2500 chromatointegrator (Hitachi) with Wakopak Eco ODS column (5 μm, 4.6 mm × 150 mm) (Fujifilm Wako Pure Chemical). The eluent was monitored at 220 nm. The mobile phase was 40% methanol/11% acetonitrile/10 mM potassium phosphate buffer (pH 7.4). The flow rate was 1.0 mL/min. The column temperature was set at 35 °C. The quantification of α- and 4-hydroxytriazolam was performed by comparing the HPLC peak areas with those of authentic standards.

### Measurement of Cyp2b10 activity and plasma efavirenz and 8-hydroxyefavirenz concentration

*In vitro* efavirenz 8-hydroxylase activity was determined as follows: a typical incubation mixture (final volume of 0.2 mL) containing 100 mM potassium phosphate buffer (pH 7.4), 1 mg/mL MLM, and 20 μM efavirenz dissolved in DMSO (1% v/v final concentration) was preincubated at 37 °C for 2 min, and the reaction was initiated by the addition of a NADPH-generating system. After 60 min incubation, the reaction was terminated by the addition of 200 μL of ice-cold acetonitrile. In a preliminary study, it was confirmed that the activity was linear with respect to protein concentration (≤ 2 mg/mL) and incubation time (≤ 90 min). After removal of protein by centrifugation at 20,380 g for 5 min, a 2 μL portion of the supernatant was subjected to LC□MS/MS. To measure plasma efavirenz and 8-hydroxy efavirenz concentration, 45 μL of acetonitrile was added to 5 μL of plasma and centrifuged at 20,380 g for 5 min. Then, 10 μL of supernatant was subjected to LC□MS/MS. An LCMS-8045 (Shimadzu, Kyoto, Japan) equipped with an LC-20CE HPLC system (Shimadzu, Kyoto, Japan) was used. The column used was a Xterra Shield RP18 column (3.5 μm, 2.1 × 150 mm: Waters, Milford, MA). The flow rate was 0.2 mL/min, and the column temperature was 40 °C. Nitrogen was used as the nebulizer gas and drying gas. The operating parameters were optimized as follows: nebulizer gas flow, 3 L/min; drying gas flow, 15 L/min; desolvation line temperature, 300 °C; and heat block temperature, 400 °C. LC□MS/MS was performed with positive electrospray ionization. The mobile phase was (A) 20 mM ammonium sulfate and 0.1% formic acid and (B) acetonitrile, and the gradient conditions of mobile phase (B) were as follows: 30 to 70% (1–5 min), 70% (5–10 min), and 30% (10–12 min). Two m/z ion transitions were monitored in multiple reaction monitoring (MRM) mode: m/z and 244.0 for efavirenz and 329.0 and 258.1 for 8-hydroxyefavirenz. The quantifications of efavirenz and 8-hydroxyefavirenz were performed by comparing the peak areas with those of authentic standards.

### Statistical analyses

Statistical significance was determined by analysis of variance followed by Dunnett’s multiple comparisons test or Tukey’s method test. Correlation analyses were performed by Pearson’s product-moment method. A value of *P* < 0.05 was considered statistically significant.

## Supporting information

Proteomics Data

## Acknowledgments

This paper was supported by Mochida Memorial Foundation for Medical and Pharmaceutical Research, Platform Project for Supporting Drug Discovery and Life Science Research (Basis for Supporting Innovative Drug Discovery and Life Science Research (BINDS)) from AMED under Grant Number JP21am0101124 (support number 2430), World Premier International Research Center Initiative (WPI), MEXT, Japan, and Research Support Project for Life Science and Drug Discovery (BINDS) from AMED under Grant Number JP22ama121046 (Y.K. and K.K) and JST CREST under Grant Number JPMJCR18S4 (Y.K.). The authors would like to thank Dr. Masahiko Negishi (National Institutes of Health, USA) for providing the ShP51 cells and FLAG-PXR plasmid, Toko Kurosaki, Yukako Sumida, Masami Morimura, Kei Yoshida, Eri Kaneda, Michika Fukino, Asahi Shibahara, and Tomoko Ashiba at Tottori University for their technical assistance.

