## Supporting information for "ncBAF, a chromatin remodeler, enhances PXR-mediated transcriptional activation in the human and mouse liver"

### SI Materials and Methods

#### *Cell cultures*

To form spheroidal aggregates, ShP51 cells were cultured on 24-well micro-space cell culture plate (D7-200-100P, Kuraray, Tokyo, Japan) with 200 μm length × 200 μm length × 100 μm depth square compartments on the culture surface. After the plate was coated with 0.05% 2-methacryloyloxyethyl phosphorylcholine polymer solution, ShP51 cells were seeded at a density of 2 × 105 cells/well on the plate. The cells were cultured for 2 weeks to form spheroids. The medium was exchanged every 2 or 3 days. Additionally, spheroids were formed using the hanging drop method with the aid of Perfecta3D Hanging Drop Plates (3D Biomatrix, Ann Arbor, MI). The cells were seeded into 96-well Perfecta3D Hanging Drop Plates at a concentration of 1,000 cells/well in a droplet containing a total volume of 40 μL. The medium was changed by drawing up 14 μL of droplet and adding 20 μL of fresh medium on alternate days for 5 days.

#### *Pull-down assay*

FLAG-PXR-transfected HEK293T cells were lysed with ice-cold lysis buffer and incubated on ice for 1 hr. After centrifugation, the supernatants were incubated with the anti-FLAG antibody-Dynabeads protein G complex for 2 hr. After washing three times using wash buffer, BRD7-His or BRD9-His-transfected HEK293T cell lysates were added to the beads and incubated at 4 °C for 12 hr. After washing using the wash buffer three times, the beads were thoroughly mixed with SDS–PAGE sample buffer. The immunoprecipitant was subjected to Western blotting.

#### *Immunofluorescence staining*

Spheroids of ShP51 cells formed by the hanging drop method were fixed with 4% paraformaldehyde for 10 min, incubated with PBS containing 0.05% Triton-X 100 for 10 min to permeate the membrane and incubated with IF blocking solution [3% BSA and 0.1% Triton X-100] for 10 min to block non-specific binding sites. The cells were then incubated with anti-human PXR antibody diluted with IF blocking solution at 4 °C for 12 hr. The cells were then rinsed three times with PBS containing 0.02% Tween 20 and incubated with IR-dye680LT goat anti-mouse IgG antibody diluted with IF blocking solution. The cells were then washed three times with PBS, incubated with PBS containing 5 μg/mL Hoechst for 5 min, and rinsed again with PBS. Fluorescent images were acquired using a Dragonfly (Oxford Instruments, Oxford, UK).

#### *Evaluation of gluconeogenesis in cellulo*

To evaluate gluconeogenesis in ShP51 cells, a glucose production assay was performed according to the method reported by ref. 1. ShP51 cells were washed with PBS and incubated in Krebs-Henseleit-HEPES buffer containing 1 mM sodium pyruvate and 10 mM sodium lactate at 37 °C for 4 hr. The buffer was collected and filtered through 0.22-μm filter, and the glucose content was determined by fluorometric enzyme assay. Briefly, 50 μL of aliquot was added to 50 μL of reaction buffer [50 mM Tris-HCl (pH 8.0), 2 mM MgCl2, 1 mM dithiothreitol, 600 μM ATP, 60 μM NADP+, 0.28 U/mL hexokinase, and 0.04 U/mL G6PDH], and the mixture was incubated at 37 °C for 30 min. The glucose concentration was determined by measuring the fluorescence intensity of NADPH at excitation 345 nm and emission 485 nm.

#### *Statistical analyses*

Statistical significance was determined by analysis of variance followed by Dunnett's multiple comparisons test or Tukey's method test. Correlation analyses were performed by Pearson's product-moment method. A value of $P < 0.05$ was considered statistically significant.

**Fig. S1.**

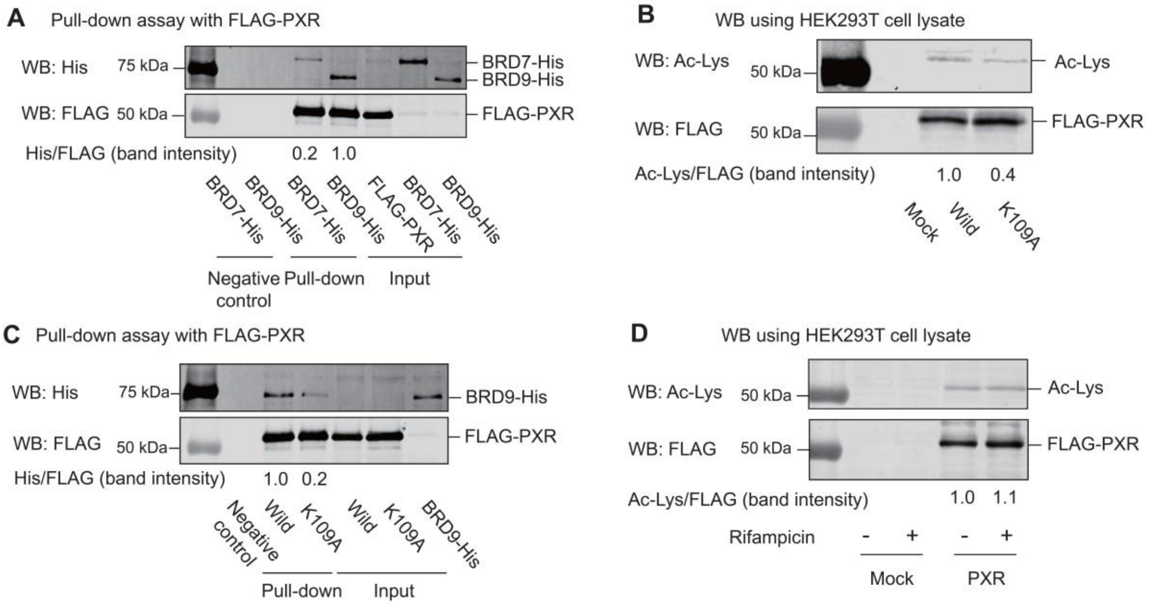


**Fig. S1.** Lysine acetylation of PXR. HEK293T cells were transfected with FLAG-PXR, BRD7-His, or BRD9-His plasmid (A), transfected with FLAG-PXR or FLAG-PXR K109A plasmid (B), transfected with FLAG-PXR, FLAG-PXR K109A, or BRD9-His plasmid (C), or transfected with FLAG-PXR or FLAG-PXR K109A plasmid followed by treatment with rifampicin for 1 hr (D). (A and C) Pull-down assay with FLAG-PXR or FLAG-PXR K109A was performed using whole lysates of HEK293T cells transfected with BRD7-His or BRD9-His, and FLAG and His tags were detected by Western blotting. (B and D) FLAG and acetylated lysine protein levels were evaluated by Western blotting. WB, Western blotting. The experiments were repeated two times with similar results.

**Fig. S2.**

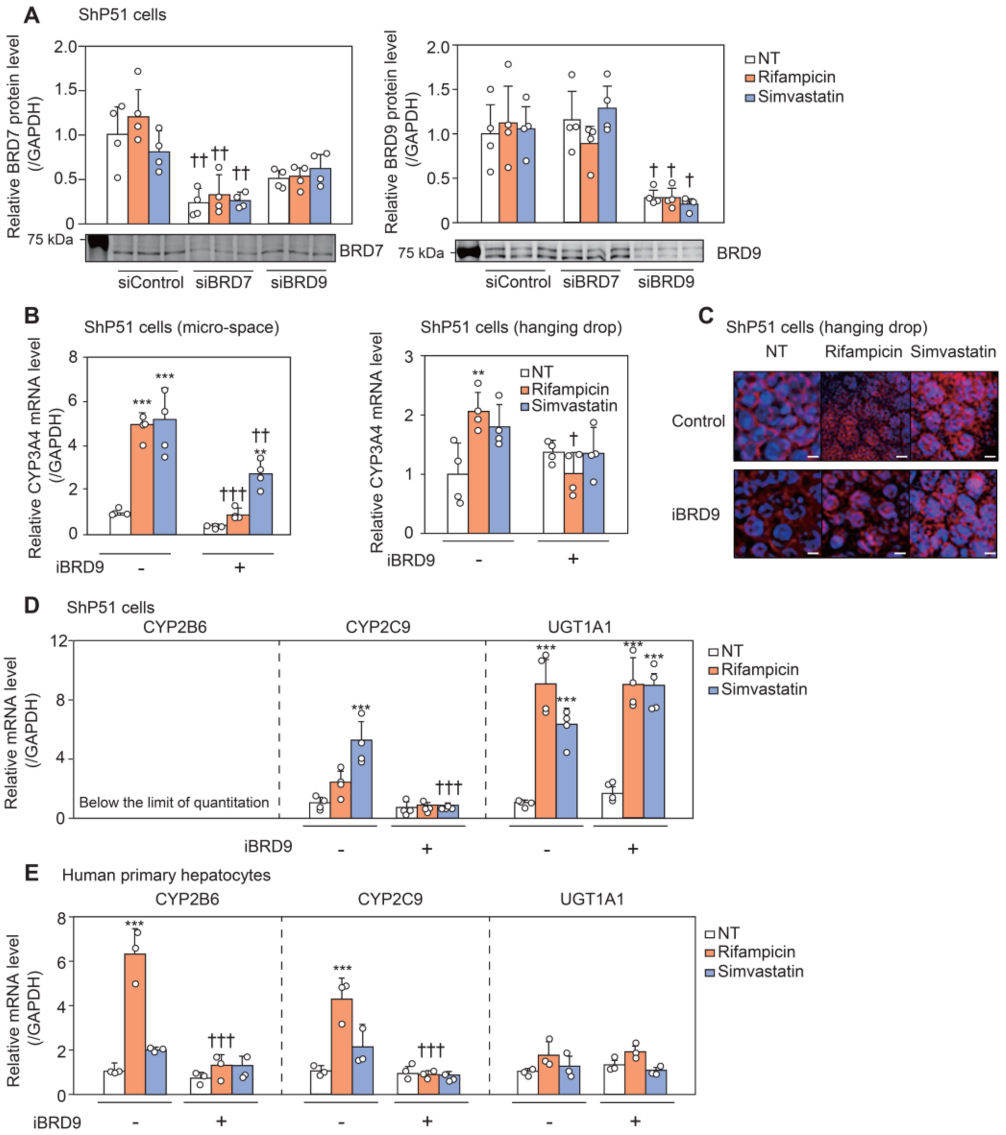


**Fig. S2.** Roles of BRD7 and BRD9 in transcriptional activation by PXR. (A) ShP51 cells were transfected with siRNA for BRD7 (siBRD7) or BRD9 (siBRD9). After incubation for 24 hr, the cells were treated with rifampicin or simvastatin. BRD7 and BRD9 protein levels were evaluated by Western blotting and normalized to GAPDH protein level. ShP51 cells in 3D culture were treated with 10 μM rifampicin or 10 μM simvastatin along with 10 μM iBRD9. (B) CYP3A4 mRNA level was evaluated by real-time RT–PCR. (C) Nuclear translocation of PXR was evaluated by immunofluorescence using anti-PXR antibody. Scale bar: 50 μm. ShP51 cells (D) and human primary hepatocytes (E) were treated with 10 μM rifampicin or 10 μM simvastatin along with 10 μM iBRD9. CYP2B6, CYP2C9, and UGT1A1 mRNA levels were evaluated by real-time RT–PCR. CYP2B6 mRNA level in ShP51 cells was below the limit of quantitation. Each column represents the mean ± SD (n = 3-4). n refers to biological repeats. ***$P < 0.001$, compared with NT, $^{\dagger}P < 0.05$, $^{\dagger\dagger}P < 0.01$, and $^{\dagger\dagger\dagger}P < 0.001$, compared with iBRD9 (-). NT: non-treatment. The experiments were repeated two times with similar results.

**Fig. S3.**

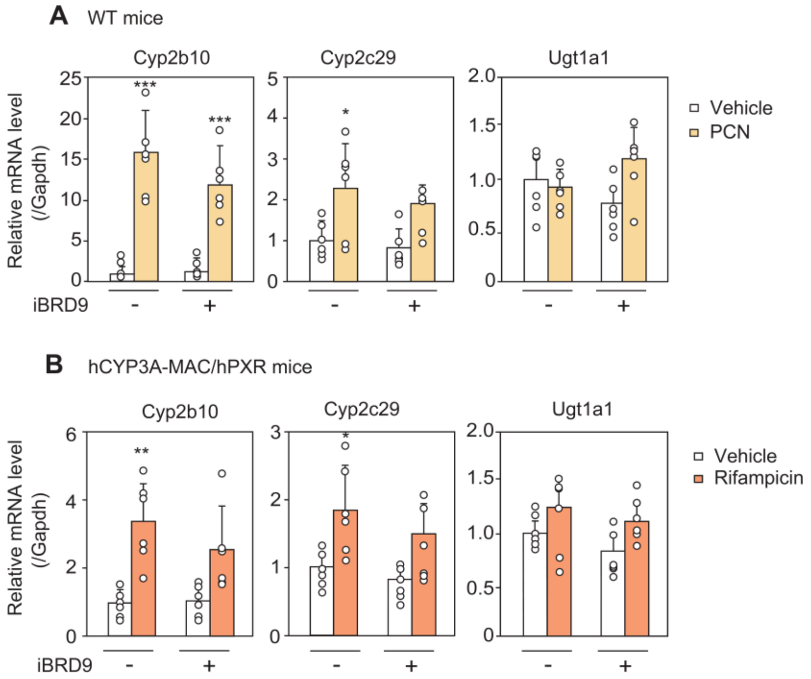


**Fig. S3.** Effects of iBRD9 on the induction of PXR downstream genes by PXR ligands *in vivo*. C57BL/6J (A) or hCYP3A-MAC/hPXR (B) male mice (n = 6) were intraperitoneally treated with 50 mg/kg PCN or 10 mg/kg rifampicin for four consecutive days and intraperitoneally treated with 10 mg/kg iBRD9 every other day. Cyp2b10, Cyp2c29, and Ugt1a1 mRNA levels were evaluated by real-time RT–PCR. Each column represents the mean ± SD (n = 6). n refers to biological repeats. **P* < 0.05, ***P* < 0.01, and ****P* < 0.001, compared with vehicle. The experiments were repeated two times with similar results.

**Fig. S4.**

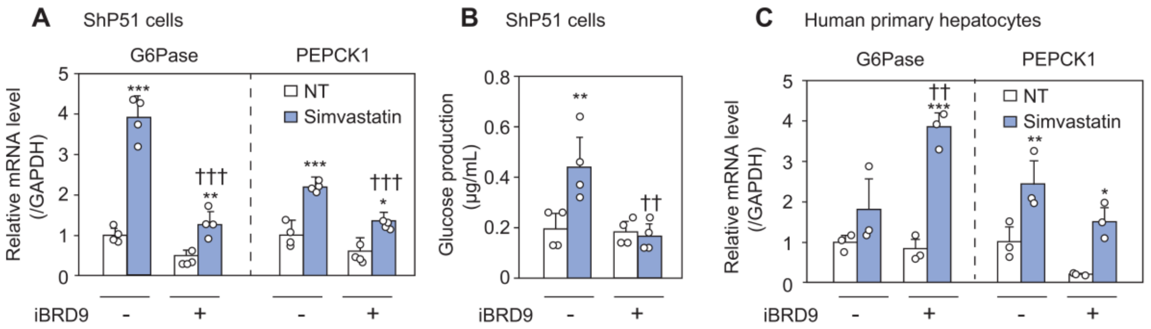


**Fig. S4.** Effects of knockdown of BRD7 or BRD9 or inhibition of BRD9 on PXR-mediated induction of gluconeogenesis. ShP51 cells (A, B) and human primary hepatocytes (C) were treated with simvastatin along with iBRD9. G6Pase and PEPCK1 mRNA levels were evaluated by real-time RT–PCR (A, C), and glucose production was evaluated (B). Each column represents the mean ± SD (n = 3-4). n refers to biological repeats. *$P < 0.05$, **$P < 0.01$, and ***$P < 0.001$, compared with NT, ††$P < 0.01$ and †††$P < 0.001$, compared with siControl or iBRD9 (-). NT: non-treatment. The experiments were repeated two times with similar results.

**Fig. S5.**

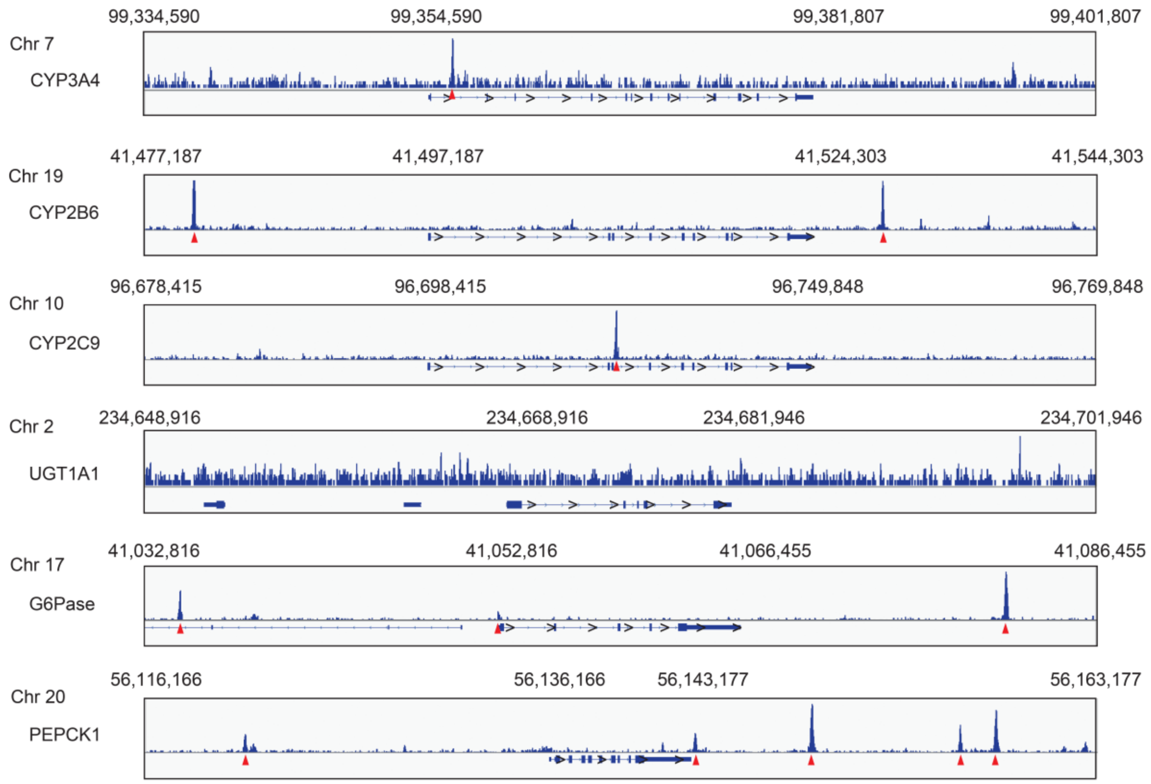


**Fig. S5.** The binding site of CTCF to PXR target genes. ChIP-seq data (SRX100531) for CTCF at *CYP3A4, CYP2B6*, *CYP2C9*, *UGT1A1*, *GSTA1*, *G6Pase*, and *PEPCK1* loci in HepG2 cells registered in the ChIP-Atlas database were visualized by Integrative Genome Viewer. Red arrows show CTCF binding sites.

**Table S1.** Sequence of primers for real-time PCR.

| Target | Primer | Sequence (5'→3') |
|---|---|---|
| hCYP3A4 | S | CAA GCT ATG CTC TTC ACC G |
| | AS | TGC AGT TTC TGG GTC CAC |
| hCYP2B6[a] | S | CTT GCG GGG ATA TGG TGT GA |
| | AS | TCA AAC AGC TGG CCG AAT AC |
| hCYP2C9[b] | S | CAG ATC TGC AAT AAT TTT TCT C |
| | AS | CTT TCA ATA GTA AAT TCA GAT G |
| hUGT1A1[c] | S | CCT TGC CTC AGA ATT CCT TC |
| | AS | ATT GAT CCC AAA GAG AAA ACC AC |
| hGSTA1 | S | GAA TTC AGT TGT CGA GCC AGG |
| | AS | TTC TGC CCC GTG CAT TGA |
| hG6Pase | S | AGC AGG TGT ATA CTA CGT GAT GG |
| | AS | CAG CAA TGC CTG ACA GGA CTC |
| hPEPCK1[d] | S | AGC TCG GTC GCT GGA TGT CAG AG |
| | AS | GTA GGG TGA ATCCGTCAG CTC GAT G |
| hGAPDH[e] | S | CCA GGG CTG CTT TTA ACT C |
| | AS | TGG GTG GCA GTG ATG GCA TGG A |
| mCyp3a11 | S | TAT ATC CCC AAA GGG TCA ACA |
| | AS | GAA GGA GAA GTT CTG CAT AAT |
| mCyp3a25 | S | CTT CAC TGT CCA GCC TTG TGA A |
| | AS | AAT TGG TTC CCT GCT GAT CTT C |
| mCyp2b10 | S | CAG ATC TTA TAC CTA TTG GAG TG |
| | AS | CAA ATG CGC TTT CCT GT |
| mCyp2c29[f] | S | GGG CTC AAA GCC TAC TGT CAT |
| | AS | GGT CAT GAG TGT AAA TCG TCT C |
| mUgt1a1 | S | GTC ACT TGC CAC TGA AAT C |
| | AS | CCA GAG GCG TTG ACA TAG |
| mGsta1 | S | TGA TGG GAT TGA AGC TGG C |
| | AS | CCA GAG GCG TTG ACA TAG |
| mSqle[g] | S | ATA AGA AAT GCG GGG ATG TCA C |
| | AS | ATA TCC GAG AAG GCA GCG AAC |
| mCd36[g] | S | CAG TCG GAG ACA TGC T |
| | AS | CTC GGG GTC CTG AGT T |
| mGapdh[h] | S | AAA TGG GGT GAG GCC GGT |
| | AS | ATT GCT GAC AAT CTT GAG TGA |
| Proximal promoter | S | AGC TCG GTC GCT GGA TGT CAG AG |
| of *hCYP3A4* | AS | GTA GGG TGA ATC CGT CAG CTC GAT G |
| Distal enhancer | S | AGC AGG TGT ATA CTA CGT GAT GG |
| of *hCYP3A4* | AS | CAG CAA TGC CTG ACA GGA CTC |
| Far enhancer | S | GTG TCA TCA CTG CTG GTA T |
| of *hCYP3A4* | AS | CAA ATC TGT TGT TTA CTC ACG TGA |

[a]Nozaki et al., 2019, [b]Katoh et al., 2004, [c]Izukawa et al., 2009, [d]Takagi et al., 2010, [e]Tsuchiya et al., 2004, [f]Kobayashi et al., 2012, [g]Gwag et al., 2019, [h]Kobayashi et al., 2010

**Table S2.** Conditions of annealing and extension for real-time RT-PCR.

| Target | Annealing | | Extension | |
|---|---|---|---|---|
| | Temperature (°C) | Time (sec) | Temperature (°C) | Time (sec) |
| hCYP2B6 | 57 | 30 | 72 | 20 |

| Target | Annealing and extension | |
|---|---|---|
| | Temperature (°C) | Time (sec) |
| hCYP3A4 | 57 | 30 |
| hCYP2C9 | 58 | 30 |
| hUGT1A1 | 64 | 20 |
| hG6Pase | 64 | 30 |
| hPEPCK1 | 66 | 20 |
| hGAPDH | 64 | 20 |
| mCyp3a11 | 57 | 20 |
| mCyp3a25 | 68 | 20 |
| mCyp2b10 | 55 | 30 |
| mCyp2c29 | 60 | 20 |
| mUgt1a1 | 55 | 20 |
| mSqle | 59 | 30 |
| mCd36 | 55 | 30 |
| mGapdh | 64 | 20 |
| Proximal promoter of *hCYP3A4* | 61 | 30 |
| Distal enhancer of *hCYP3A4* | 52 | 30 |
| Far enhancer of *hCYP3A4* | 55 | 20 |

**Dataset S1 (separate file).** Proteomics dataset is included in the SI appendix.
